# Performance of semi-automated hippocampal subfield segmentation methods across ages in a pediatric sample

**DOI:** 10.1101/064303

**Authors:** Margaret L. Schlichting, Michael L. Mack, Katharine F. Guarino, Alison R. Preston

**Affiliations:** Center for Learning and Memory, The University of Texas at Austin; Departments of Psychology, The University of Texas at Austin; Departments of Neuroscience, The University of Texas at Austin; Department of Psychology, University of Toronto

**Keywords:** Development, structural MRI, volume, high-resolution MRI, reliability

## Abstract

Episodic memory function has been shown to depend critically on the hippocampus. This region is made up of a number of subfields, which differ in both cytoarchitectural features and functional roles in the mature brain. Recent neuroimaging work in children and adolescents has suggested that these regions may undergo different developmental trajectories—a fact that has important implications for how we think about learning and memory processes in these populations. Despite the growing research interest in hippocampal structure and function at the subfield level in healthy young adults, comparatively fewer studies have been carried out looking at subfield development. One barrier to studying these questions has been that manual segmentation of hippocampal subfields—considered by many to be the best available approach for defining these regions—is laborious and can be infeasible for large cross-sectional or longitudinal studies of cognitive development. Moreover, manual segmentation requires some subjectivity and is not impervious to bias or error. In a developmental sample of individuals spanning 6-30 years, we assessed the degree to which two semi-automated segmentation approaches—one approach based on Automated Segmentation of Hippocampal Subfields (ASHS) and another utilizing Advanced Normalization Tools (ANTs)—approximated manual subfield delineation on each individual by a single expert rater. Our main question was whether performance varied as a function of age group. Across several quantitative metrics, we found negligible differences in subfield validity across the child, adolescent, and adult age groups, suggesting that these methods can be reliably applied to developmental studies. We conclude that ASHS outperforms ANTs overall and is thus preferable for analyses carried out in individual subject space. However, we underscore that ANTs is also acceptable and may be well-suited for analyses requiring normalization to a single group template (e.g., voxelwise analyses across a wide age range). Previous work has supported the use of such methods in healthy young adults, as well as several special populations such as older adults and those suffering from mild cognitive impairment. Our results extend these previous findings to show that ASHS and ANTs can also be used in pediatric populations as young as six.

---

Recent years have seen increasing research interest in how the hippocampus (HPC) develops, both in terms of structure and function. In particular, work combining high-resolution structural imaging methods with new analysis techniques (Gogtay et al., 2006; Lee et al., 2014; Lin et al., 2013) has suggested that the HPC may continue to change in subtle ways through at least late childhood, and perhaps even into early adulthood. For instance, different developmental trajectories have been observed across the anterior-posterior axis of the hippocampus, with anterior regions generally showing decreases and posterior regions showing increases in size with age (Demaster et al., 2013; Gogtay et al., 2006). Theoretical and animal models suggest that anatomical pathways within the hippocampal circuit may also mature at different rates (Gómez and Edgin, 2015; Lavenex and Banta Lavenex, 2013), which could give rise to the different developmental trajectories sometimes observed across subfields (Daugherty et al., 2016; Tamnes et al., 2014a): the *cornu ammonis* (CA) fields, dentate gyrus, and subiculum. These substructures within HPC have unique anatomical and functional characteristics in the mature brain (Carr et al., 2010; Manns and Eichenbaum, 2006). Thus, these perspectives suggesting variability in the developmental trajectory of different HPC substructures make a host of predictions about when the functions of each region—and thus, the corresponding mnemonic behaviors—each reach maturity. A spate of recent work has jumpstarted the enterprise of empirically testing these hypotheses (Canada et al., 2018; Daugherty et al., 2017; Keresztes et al., 2017; Ngo et al., 2017; Riggins et al., 2018, 2015; Schlichting et al., 2017; Tamnes et al., 2018, 2014b).

Answering questions about HPC development requires the application of advanced neuroimaging techniques to pediatric populations. For instance, future studies will likely seek to use high-resolution functional MRI (fMRI) to interrogate activation profiles within different subfields of the hippocampus and characterize how they change over development. This approach requires not only the acquisition of high-resolution fMRI data, but also the ability to reliably demarcate HPC subfields as anatomical regions of interest (ROIs) and/or normalize individual anatomical images to a custom template generated for the purpose of localizing activations to particular HPC subfields. These methods should be able to be carried out in a consistent manner on participants spanning a wide age range—preferably in a manner that is easily reproducible across studies and research groups.

There is an increasing push toward larger sample sizes in neuroscience research (Button et al., 2013), and developmental work is no exception. Developmental questions regularly require large sample sizes due to the defining characteristic of the discipline as one interested in the impact of age, either as an individual difference in a cross-sectional study or in a longitudinal design. For example, developmental researchers might investigate how brain-behavior relationships differ as a function of age or characterize within-participant change by acquiring data at multiple timepoints in a longitudinal study. As manual delineation of HPC subfields is time-consuming to both master and perform, the large amount of data required by many developmental researchers renders this best available method of hippocampal subfield segmentation intractable for many researcher groups. Moreover, as this process involves consideration of both probabilistic boundaries as well as those identifiable from visible landmarks, there is a degree of subjectivity that seeps its way into segmentations produced manually.

The questions that remain outstanding in the literature—as well as the practical concern of too much data, too little time—motivated us to assess how semi-automated HPC subfield segmentation methods trained on a particular tracing protocol compare with segmentations manually delineated using that same protocol. Moreover, we were especially interested in characterizing the performance of these approaches in younger participants. The majority of existing studies investigating the development of hippocampal subfields with automated segmentation methods (Krogsrud et al., 2014; Tamnes et al., 2014a) have used Freesurfer (http://surfer.nmr.mgh.harvard.edu/) and provide no quantitative assessment of the method. Importantly, these studies used “routine” T1-weighted MR images (borrowing terminology from Yushkevich et al., 2015b) of relatively low resolution (~1mm isotropic voxels) on which the internal structure of the HPC is not visible. Here, we use “dedicated” high-resolution (0.4 × 0.4 × 1.5 mm) T2-weighted images with a very specific orientation—perpendicular to the HPC long axis—that enable us to visualize HPC subfields and extend on this prior work.

In terms of systematic assessments of automated methods over development, one study (Schoemaker et al., 2016) did investigate the correspondence between manual tracing of the *overall* HPC in children and two automated segmentation methods: Freesurfer (Desikan et al., 2006; http://surfer.nmr.mgh.harvard.edu/) and FIRST, part of FSL (Smith et al., 2004; www.fmrib.ox.ac.uk/fsl). Both automated methods failed to reach acceptable levels of reliability; however, it was unclear whether the failure of these automated methods was due to the developmental stage of the sample or is due to a more general concern about these tools, as similarly low reliability has been reported previously in adults (Doring et al., 2011; Pardoe et al., 2009). However, other groups have reported success with automated methods for segmenting the overall HPC in preterm neonates (Guo et al., 2015) and toddlers (Gousias et al., 2008). Thus, there is mixed support in the literature for using automated HPC segmentation methods in developmental work.

With regards to automated subfield segmentation, while there is no dearth of literature validating the use of such methods in other special populations such as older adults and individuals suffering from mild cognitive impairment, Alzheimer’s Disease, and psychoses (Pipitone et al., 2014; Yushkevich et al., 2010; Paul A. Yushkevich et al., 2015b), there is only one paper to our knowledge that formally assessed automated segmentation methods in a pediatric sample (Bender et al., 2018). That study found acceptable levels of performance for one of the automated methods we test in the present paper (ASHS) for an early lifespan sample aged 6-26 years, and when restricted to the HPC body. However, as the main goal of that work was to compare across multiple atlases applied to the early lifespan sample rather than segmentation performance across specific ages, it remains unknown whether the performance of automated methods varies across different developmental periods. Given that the primary goal of much work in developmental cognitive neuroscience is to directly compare individuals at different stages of development—for example, *how do children differ from adults in X structure or Y function?*—it remains a critical open question whether such methods show similar performance across age groups or worse performance in children and adolescents.

In the present study, we sought to assess how comparable two semi-automated subfield segmentation techniques are to manual subfield delineation in a pediatric sample. Importantly, these segmentations are all derived from a single tracing protocol; manual delineation was carried out by a single rater referencing histological work and printed atlases of human hippocampus (Duvernoy, 1998; Mai et al., 2007; West and Gundersen, 1990). Selecting and developing an in-house tracing protocol is common among research groups studying subfield function using fMRI in humans. We thus chose analyses that would be informative in the context of this workflow—wherein a tracing protocol may be pre-determined but the semi-automated segmentation procedures are flexible. We chose to have our hand-delineated regions ‘drawn’ by a single rater to maximize the consistency of regions both (1) going into ‘training’ the semi-automated methods as well as (2) being combined across participants to yield the ultimate validity metrics. Our aim is focused not on validating this manual tracing protocol, but to systematically evaluate how well semi-automated segmentation methods can approximate the same manual tracings in various age groups.

For our semi-automated methods, we selected two approaches with which we could use our atlas of choice, allowing us to generate regions that would be directly comparable to our drawn ROIs (**Fig. 1**): reverse normalization of regions drawn on a custom template to each participant’s native space using Advanced Normalization Tools (ANTs; Avants et al., 2011), and automated segmentation on each participant’s brain implemented using the Automated Segmentation of Hippocampal Subfields (ASHS) software (Paul A. Yushkevich et al., 2015b). Critically, both methods also allowed us to use the same reference (i.e., ANTs template and ASHS atlas) and analysis strategy across age groups, which—assuming there is no systematic, age-related bias—is important for making direct comparisons across groups. We then compared the ROIs generated by each automated method to the manually drawn regions separately for participants in child, adolescent, and adult age groups to test for possible differences in the convergent validity of these approaches across our age range of interest. Results revealed little evidence that the validity of ANTs and ASHS varied as a function of age group; those differences that did exist were numerically small. While both methods performed well overall, ASHS outperformed ANTs on several of our metrics. For researchers wishing to employ our segmentation approach on their own datasets, our ANTs template and ASHS atlas have been made available for download: https://osf.io/hrv9n/.

**Figure 1.**
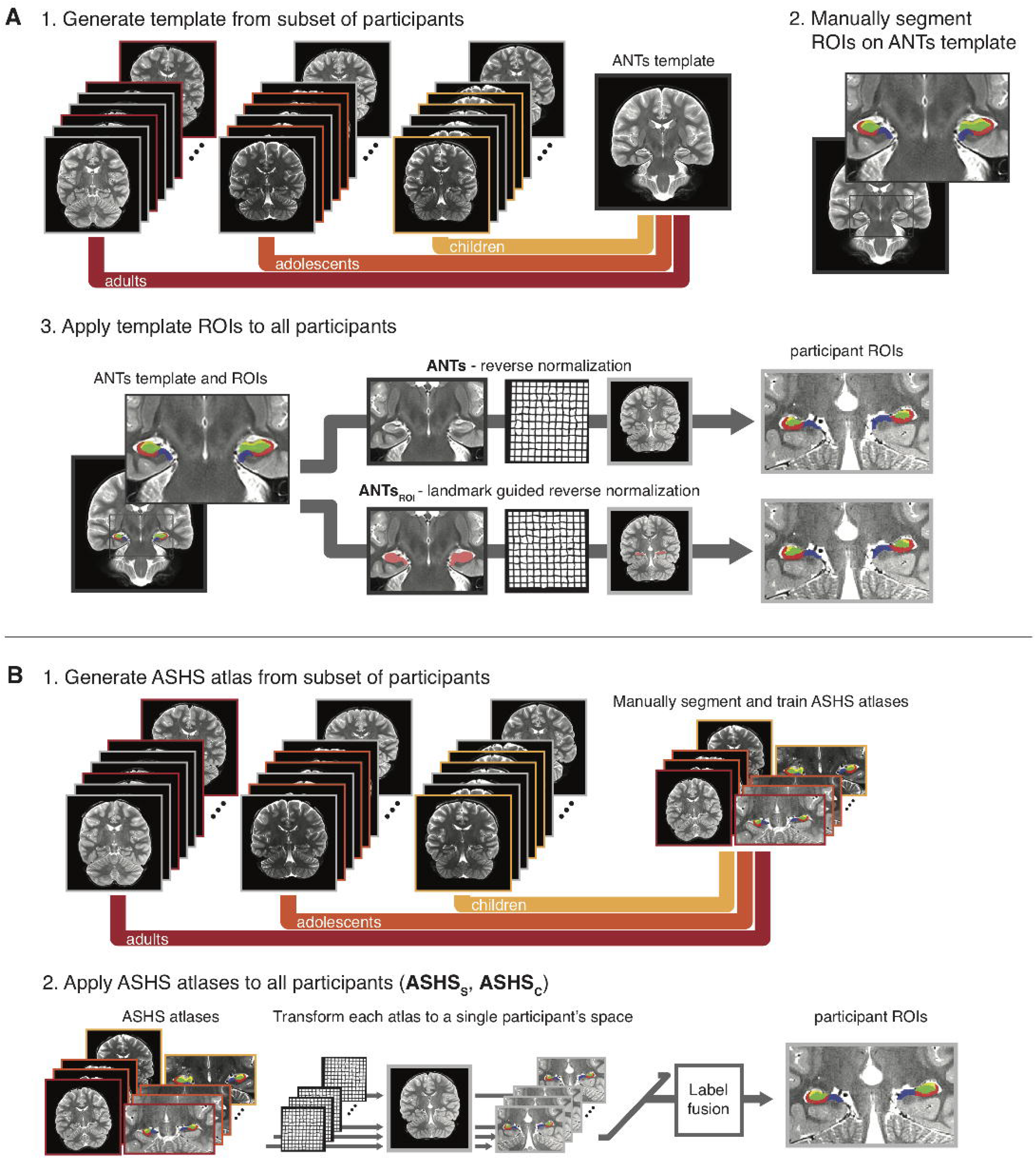
Schematic depiction of methods compared. **A)** ANTs segmentation methods. **1.** Subsets of participants were selected from the three age groups to generate a single ANTs template. **2.** Then, hippocampal subfield ROIs were manually delineated on the ANTs template. **3.** Finally, the ANTs template ROIs were reverse normalized to each participant’s native space. For the **ANTs** method (top path), a nonlinear warp was estimated based uniformly on the entire ANTs template and participant anatomical volumes. For the **ANTs_ROI_** method (bottom path), the nonlinear warp was estimated through landmark matching of the whole hippocampus. For the main analysis, both ANTs-based methods were performed for all participants to create two sets of participant-specific subfield ROIs. **B)** ASHS segmentation methods. **1.** Subsets of participants from the three age groups were selected for training custom ASHS atlases. The ASHS training procedure was run using the manually delineated subfield segmentations for the atlas participants. **2.** Then, the ASHS atlases were applied to all participants. Briefly, the ROIs from each atlas participant were all nonlinearly warped to a single participant. Then, a label fusion procedure combined the transformed atlas segmentations to generate final, participant-specific ROIs. In the main analysis, atlas participants’ own segmentations were excluded from the label fusion procedure to reduce bias in their final segmentation. The whole procedure was performed twice: once with DG and CA_2,3_ separated (**ASHS_S_**), and once with them combined (**ASHS_C_**). Figure depicts the main analysis including all participants; we additionally performed follow-up analysis omitting participants who went into the generation of the ANTs template or ASHS atlas.

## MATERIALS AND METHODS

### Participants

Ninety volunteers participated in the experiment across child (ages 6-11 y; N=31), adolescent (12-17 y; N=25), and adult (18-30 y; N=34) age groups. Participants of all ages were recruited from the greater Austin area and were thus a mixture of University-affiliated and community individuals. The consent/assent process was carried out using age-appropriate language in accordance with an experimental protocol approved by the Institutional Review Board at the University of Texas at Austin. For participants under 18, assent was obtained from the participant and permission was obtained from his/her parent or guardian. Adults provided consent. All participants received monetary compensation and a small prize for their involvement in the study.

Participants were screened for psychiatric conditions using the Child Behavior Checklist (CBCL; completed by the parent/guardian of participants aged 6-17; Achenbach, 1991) and the Symptom Checklist 90-Revised (SCL-90-R; adults; Derrogatis, 1977). IQ was assessed using the Wechsler Abbreviated Scale of Intelligence, Second Edition (WASI-II; Wechsler, 1999). The intelligence measure of interest was the full-scale IQ composite score (FSIQ-2), which includes vocabulary and matrix reasoning subtests.

From the original group of 90 participants, individuals were excluded from subsequent analysis if they met any of the following criteria: (1) CBCL score in the clinical range (N=1 child; N=1 adolescent) or SCL-90-R score greater than 1 SD above the mean of a normative sample (N=9 adults); (2) presence of a psychiatric condition (N=1 adult); (3) did not complete the MRI portion (N=3 children; N=4 adults); (4) handedness concerns (N=1 adolescent); (5) MRI data not of acceptable quality (N=4 children); or (6) automated segmentation failed (N=1 adolescent; N=2 adults). Samples of images of unacceptable quality and for which automated segmentation failed are provided in **Online Supplementary Figure 1**. Participants were excluded for automated segmentation failure when visual inspection revealed that the hippocampus as a whole was grossly mislabeled (e.g., with HPC extending through white matter and into medial temporal lobe cortex). For two of these participants (adults), both ASHS and ANTs failed; only ANTs failed for the third participant (adolescent). No participants scored below our inclusion threshold for IQ (> 2 SD below the mean).

Through the revision process, we additionally identified two outlier participants using a Mahalanobis distance test of cross-method correspondence of overall HPC volumes. Mahalanobis distance provides a measure of the likelihood of a data point in a multivariate dataset given a dataset’s covariance structure. Potential outliers were identified by comparing each participant’s bilateral HPC volume as defined by manual tracing with each semi-automated method. A threshold of χ^2^_2,.99_ flagged one child and one adolescent participant as potential outliers (i.e., showing lower correspondence between methods than would be typical for their age group). Upon manual inspection, these individuals were determined to have poor manual tracing and were subsequently removed from all analyses. The final sample included a total of 62 right-handed participants (22 children, 13 females, ages: 6.08-11.83 y, 9.52 ± 0.38 y, FSIQ-2: 84-142, 118.86 ± 2.90; 21 adolescents, 10 females, ages: 12.08-17.33 y, 14.26 ± 0.37 y, FSIQ-2: 92-130, 110.95 ± 2.58; 19 adults, 10 females, ages: 18.67-28.92 y, 23.86 ± 0.78 y, FSIQ-2: 92-129, 113.00 ± 2.76).

### Experiment Overview

The experiment comprised two visits. On the first visit, participants were exposed to the MRI environment using a mock scanner, completed paper-based screening measures (CBCL or SCL-90-R; WASI-II), and performed a battery of cognitive tasks (not discussed here). MRI scanning took place during the second visit.

### MR Data Acquisition

Imaging data were acquired on a 3.0T Siemens Skyra MRI. Two to three oblique coronal T2-weighted structural images were acquired perpendicular to the main axis of the HPC (TR=13150 ms, TE=82 ms, 512 x 60 x 512 matrix, 0.4 x 0.4 mm in-plane resolution, 1.5 mm thru-plane resolution, 60 slices, no gap, acquisition time 6:36). Coronal images of acceptable quality as determined by visual inspection (e.g., absence of motion artifacts that would prevent visualization of the hippocampal sulcus) were coregistered using ANTs (Avants et al., 2011) and averaged to improve visualization of the internal structure of the hippocampus, yielding a single mean coronal image per participant. A T1-weighted 3D MPRAGE volume (256 x 256 x 192 matrix, 1 mm^3^ voxels) was also collected.

### Baseline method: Manual hippocampal subfield delineation

We used manual demarcation to define subfields on each participant’s anatomical image (hereafter denoted Manual), which is typically considered the most accurate method for anatomical volume assessment (Rodionov et al., 2009). As such, all semi-automated methods described were compared to Manual as a reference in a pairwise fashion. HPC regions of interest (ROIs) were delineated on each participant’s mean coronal image by a single rater (KFG) following a published protocol (Bonnici et al., 2012) and further referencing printed hippocampal atlases and histological work (Duvernoy, 1998; West and Gundersen, 1990). We chose to have a single rater perform the segmentations to reduce the variance across manual tracings associated with rater differences. However, we underscore that in general a multi-rater approach in which the tracing protocols are validated through high agreement with *other* expert raters would be ideal (see Limitations section of Discussion for more on this point). The rater was blind to participant identity (including but not limited to age, sex, and performance on behavioral tasks), and images were cropped to obscure overall head size to ensure unbiased application of the segmentation protocol across participants of different ages.

We followed the segmentation protocol described by Bonnici and colleagues (Bonnici et al., 2012) referencing additional atlases and histological work (Duvernoy, 1998; Mai et al., 2007; West and Gundersen, 1990). HPC was segmented into the following subfields: *cornu ammonis* fields 1 (CA_1_) and 2/3 (combined; CA_2,3_), dentate gyrus (DG), and subiculum. Segmentation was performed across the entire extent of the HPC long axis, with the exception of the most posterior slices on which subfields could not be reliably delineated. For this region, we created a combined posterior HPC ROI (as was done in Yushkevich et al., 2015b). All subfields and posterior HPC were summed to create overall HPC ROIs. As functional neuroimaging studies often interrogate activation within a combined DG/CA_2,3_ region, we also summed individual DG and CA_2,3_ regions to create DG/CA_2,3_. While our main analyses used bilateral regions of interest, for completeness we also report all metrics split by hemisphere (see Inline Supplementary Information).

Furthermore, it is possible that validity of the semi-automated methods may be lower in the HPC head, as the boundaries are more complex in shape and have fewer visible landmarks than those in the HPC body. Therefore, in addition to the overall HPC subfields described above, we also investigated convergent validity for subfields restricted to the head versus those in the body (see Supplementary Information for results). The posterior boundary of the HPC head was the last slice on which the uncal apex was visible (Poppenk and Moscovitch, 2011; Weiss et al., 2005). The posterior boundary of the HPC body was one slice anterior to the first slice showing separation of the fornix from the HPC (Watson et al., 1992).

We quantified the reliability our manual segmentation approach by computing intra-class correlation (ICC(2,1) absolute agreement of single measures; Shrout and Fleiss, 1979) and spatial overlap (Dice similarity coefficient [DSC]; Dice, 1945) for a subset of our participants. As all of our manually delineated regions were drawn by a single rater, intra-rater reliability—in other words, the correspondence of volumes drawn by the same rater across tracing occasions—is the most relevant to the present study (for a similar approach, see Yushkevich et al., 2015b). For this purpose, our rater (KFG) manually delineated subfields for all child participants for a second time at a delay of at least one year. We focused quantification of intra-rater reliability on the child group because we reasoned this group might be the most anatomically variable and/or have the lowest image quality, and therefore might be the most difficult to segment. ICC was at acceptable levels (≥0.80) for a majority of the subfields that are the focus of the present manuscript (HPC: 0.82, CA_1_: 0.81, DG: 0.83, DG/CA_2,3_: 0.87); however, two regions fell below standards in ICC for manual demarcation (CA_2,3_: 0.66, SUB: 0.68). Spatial overlap (DSC) was high for the majority of regions (HPC: 0.91, CA_1_: 0.77, SUB: 0.76, DG: 0.78, DG/CA_2,3_: 0.84) with the exception of one region (CA_2,3_: 0.50). The majority of these results—overall HPC, CA_1_, DG, and DG/CA_2,3_—are in the same range as previous reports for manual demarcation of hippocampal subfields (Lee et al., 2014; Mueller et al., 2007; Wisse et al., 2012; Yushkevich et al., 2010) and suggest sufficiently reliable segmentation within rater across time. However, in order to provide a stringent baseline for evaluating the semi-automated segmentation methods, we limit our interpretation of validity measures to only those regions showing acceptable intra-rater reliability (ICC≥0.80). For completeness, we include results for CA_2,3_ and SUB in the subsequent sections, but caution against interpreting these findings as evidence in support of one method over another.

### Automated methods for comparison

#### Comparison method 1: Custom template ROIs reverse normalized (ANTs)

For the first comparison method (**Fig. 1A**), we defined ROIs on a template brain image, and then back-projected the regions into each participant’s native space. We first generated a series of custom templates from the mean coronal images of a subset of participants with canonical hippocampi. “Canonical” was defined subjectively, taking into account features of both the individual’s neuroanatomy (e.g., morphometry of the hippocampus) as well as the MR acquisition (e.g., whether landmarks could be readily visualized given slice orientation and image quality). Template generation and normalization were carried out using ANTs version 2.1.0 (Avants et al., 2011). Multiple templates were generated using different subsets of participants. We then selected the best group template, i.e., one that was free from artifacts and for which the HPC subfield landmarks could be visualized. Efforts were made to include participants who spanned our age range in the generation of all templates. The final chosen template used for both comparison methods 1 and 2 was created from the mean coronal images of 10 individuals (overall age range: 7.5-28.42 y; N=3 in child age range: 3 females, mean 9.3 ± SEM 1.3 y; N=3 in adolescent age range: 1 female, 14.31 ± 1.11 y; N=4 in adult age range: 2 females, 23.0 ± 2.31 y). The number 10 was chosen in accordance with the recommended guidelines for optimal template construction in ANTs, which note that the final outcome (i.e., the resulting template image) stabilizes across subsets of participants when approximately 10 images are used as input (see ANTs documentation, available at the following URL as of the writing of this paper: http://stnava.github.io/ANTs/).

The manual rater (KFG) then segmented the HPC on the group template into subfields using the protocol described above for the manual ROI delineation. In other words, the subfields were demarcated on the group coronal template in the same way as they were for an individual brain. We then back-projected all ROIs into the native space of each participant as follows. First, we computed the nonlinear transformation (which in the present paper always includes an affine step to initialize the registration) from the individual’s mean coronal to the group template using the following settings: image metric: probabilistic; transformation: symmetric normalization; regularization: Gaussian. We then applied the inverse transformation to each ROI.

This method does require some time and expertise on the part of the researcher to implement. First, it may be the case that multiple templates need to be generated to yield one that will allow for manual segmentation and satisfactory normalization. Second, the researcher must manually segment the regions on the template according to their desired protocol. Thus, this procedure is not fully automated; however, it does significantly cut down the hands-on time required by the researcher relative to manual segmentation of each individual, making it tractable for larger N studies.

#### Comparison method 2: Custom template ROIs reverse normalized using ROI-guided methods (ANTs_ROI_)

The second comparison method used a procedure identical to the one described for ANTs above, with the single exception that regions were back-projected into native space using nonlinear transformations computed with ROI-guided methods implemented using landmarkmatch, part of the ANTs toolbox (**Fig. 1A**). Hereafter, this method of ROI-guided ANTs normalization is denoted ANTs_ROI_. Specifically, the Manual HPC for each individual was used to guide the normalization to the custom group template, with the Manual HPC template ROI serving as the target (weight: 1). The inverse of the computed transformation was then applied to the group template ROIs, such that regions were back projected to native space.

As with ANTs, ANTs_ROI_ requires template selection and manual demarcation on the part of the researcher. In addition, a significant downside to ANTs_ROI_ as compared with ANTs is that implementing this approach requires a rater to trace HPC on each participant’s brain individually. This practical concern should be taken into consideration when choosing a segmentation method.

#### Comparison methods 3 and 4: ASHS automated segmentation using custom atlases

For the third and fourth comparison methods (both depicted in **Fig. 1B**), we built custom atlases for use with the Automated Segmentation of Hippocampal Subfields (ASHS) software (version 0.1.0, rev 103 downloaded on 4/14/2016) (Paul A. Yushkevich et al., 2015b). ASHS is an open-source software package that uses both T1- and T2-weighted images to automatically segment the medial temporal lobe into subregions. It can be used with an included atlas, or retrained to use any segmentation protocol chosen by the user. For the present study, we generated custom atlases based on the Manual ROIs for a subset of participants with canonical hippocampi. Nine participants from each of the three age groups (total N=27) were selected for atlas generation (age range: 6.08-28.75 y, hereafter termed “atlas participants”; N=9 in child age range: 6 females, 9.18 ± 0.61 y; N=9 in adolescent age range, 6 females, 14.99 ± 0.58 y; N=9 in adult age range: 5 females, 23.89 ± 1.21 y). This number was chosen to be in the 20-30 range recommended in the ASHS software documentation (https://sites.google.com/site/hipposubfields/building-an-atlas). We then built two atlases: one that combined across DG and CA_2,3_ (ASHS combined, hereafter ASHSC), and one that included them as separate regions (ASHS separated, hereafter ASHSS). These two comparison methods were otherwise identical.

Following atlas generation, automated segmentation was carried out on all participants. The standard ASHS segmentation procedure allows all atlas participants to “vote” on the subfield label for each voxel, an approach known as multi-atlas label fusion (Paul A. Yushkevich et al., 2015b). In our case, as the atlas participants were also members of our target segmentation sample, we modified this procedure to remove the possibility of any bias in the label fusion step: Namely, for those participants whose manual ROIs went into atlas generation, their vote was excluded from subfield label fusion. This avoids a participant’s own manual segmentation potentially driving a highly accurate (but biased) segmentation. Our modified approach allows for an unbiased comparison to the ANTs methods for the purposes of the current work. Note that when ASHS is used in practice, manually drawn ROIs from atlas subjects rather than the ASHS atlas could be used for characterizing anatomical and functional measures of those subjects.

Manually delineated ROIs are needed for the initial step of generating a custom atlas. However, once an atlas has been created, it can easily be applied to an entirely new sample, which would make the segmentation entirely automated. Users wishing for a fully automated pipeline may also download and use an existing, publicly available atlas.

### Volume extraction

Each comparison method resulted in HPC, CA_1_, CA_2,3_, DG, DG/CA_2,3_, and subiculum regions in each participant’s native space. Because the most posterior portion of the hippocampus was not segmented into subfields on the majority of participants, we consider validity of subfield delineation only on slices for which subfields were drawn. Raw volumes were extracted for all methods and then adjusted for differences in overall head size across all participants as follows. Intracranial volume (ICV) was estimated from each participant’s T1-weighted image using Freesurfer (Desikan et al., 2006). We extracted volumes for each ROI and participant. To account for differences in overall head size, volumes for each ROI were adjusted for ICV using an analysis of covariance approach (Raz et al., 2005). Specifically, each ROI (including overall HPC) was regressed on ICV across the age range to determine the slope (βICV) of the relationship between overall head size and ROI volume. Raw ROI volumes were then adjusted to correct for this relationship by subtracting the product of mean-centered ICV measures and β_ICV_ from each ROI. This procedure removes the statistical relationship between ICV and ROI volumes. Adjusted rather than raw ROI volumes then went into all subsequent volume-based analyses. We reasoned that this was the best choice in this context, as the ICV-normalized values are what would ultimately go into a final analysis in any paper investigating individual differences in region size or brain-behavior relationships.

The convergent validity of the automated methods were compared across age groups using various metrics as described below. Basic quality assurance was performed to exclude participants (N=5) whose overall HPC was grossly mislabeled, as described above. This step was the only quality control we carried out for the automated segmentations; no manual editing of regions was performed.

### Comparing automated segmentation methods with manual demarcation

#### Spatial overlap (DSC)

We first interrogated how much each automated method agreed with the Manual regions in terms of their spatial overlap. Spatial overlap was indexed using Dice similarity coefficients (DSC; Dice, 1945). For region X segmented using methods A and B, DSC is defined as:

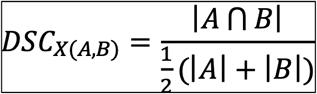

where A and B are the set of voxels marked as region X by methods A and B, respectively. DSC was computed for each participant, ROI, and comparison method; the resulting values were then averaged across participants within each age group. Note that as DSC is a measure of spatial overlap rather than volume, it is not corrected for ICV.

#### Edge agreement across methods

While DSC quantifies the amount of overall spatial overlap between two methods, it does not indicate where the disagreements among methods lie. To further characterize the spatial locations at which our automated methods agree (or disagree) with our Manual subregions, we indexed agreement at particular voxels in a mapwise fashion (see Yushkevich et al., 2015 for a similar approach). For each participant and method, we created an image indicating edge voxels (i.e., voxels on either side of the boundary between two subfields, or between a subfield and the outside of HPC). We then compared the edges for each method to the Manual edge map in native participant space, characterizing agreement as voxels that were marked as edges by both methods. These maps were then normalized to the group template for display and averaged across individuals within each age group to serve as a qualitative representation of edge agreement. The intensities of the resulting maps thus indicate the proportion of participants that showed agreement between the two methods at each specific voxel.

In addition to these qualitative results, we also quantified the degree to which these edges overlapped with those determined via Manual demarcation across the anterior-posterior HPC axis, again motivated by concerns that segmentation in anterior HPC may be less reliable. This analysis was performed in the native space of each participant (i.e., no spatial normalization was performed) using the edge maps described above. For each participant and semi-automated method, we calculated DSC overlap for the ANTs- or ASHS-generated edge map with the manually drawn edges within each HPC slice. For each participant, slice location was centered on the boundary between HPC body and head as determined by author KFG during initial manual demarcation. DSC values were averaged within age groups at each centered slice that contained manually-demarcated ROIs from at least 90% of the participants. This approach provides a measure of segmentation error specific to the edges and boundaries of ROIs at each anterior-posterior position within HPC that can be compared across age groups and methods. It is worth noting that due to the fewer number of voxels in edge maps this measure of overlap will almost necessarily lead to lower agreement than what is typically observed with volume-based DSC measures. Although an acceptable threshold of edge-based DSC has not (to our knowledge) been established, the relative comparisons between methods and age groups provide insight into the spatial agreement of the different methods. Note that both edge overlap analyses measure spatial overlap rather than volume, as such they are not corrected for ICV.

#### Volume correspondence: Intra-class correlation

We next assessed the degree of correspondence between regional volumes determined using each method, irrespective of spatial agreement. We computed intra-class correlation (ICC) to measure absolute agreement of single measures using a two-way random effects ANOVA model (i.e., ICC(2,1), Shrout and Fleiss, 1979). ICC values near 1 indicate high agreement in the volumes derived across methods, while values near 0.5 indicate poor agreement. Prior work on hippocampal subfields in developmental samples (Lee et al., 2014) has considered values exceeding 0.70 to represent an acceptable or “good” level of agreement, in line with recommendations from Bartko, 1991.

#### Bias

One possibility is that the degree to which volumes are over- or under-estimated by a given method varies as a function of the region’s size. This might be of particular concern for researchers investigating hippocampal subfields (as subfields vary substantially in size) and/or their development (as overall size will differ across age groups). To assess this possibility, we generated Bland-Altman plots (Bland and Altman, 2007), which show the difference in volume of the two methods (e.g., ANTs-Manual on the y-axis) as a function of the average of the two methods (x-axis). If the automated method is not systematically over- or under-estimating the region volumes relative to Manual, the difference values should be centered around zero. Additionally, we performed ANCOVAs to determine whether there were significant main effects of average volume, age, and/or a volume x age interaction. Significant effects of volume indicate bias in the volume estimation across all three age groups, with the size of the ROI predicting the degree to which volume is over or underestimated by the automated method. Significant effects of age indicate that the degree of volume under- or overestimation differs across the three age groups. A significant volume x age interaction would indicate that the bias profile differs across the age groups.

### Assessing reliability within segmentation method

#### Volume correspondence within an individual: inter-hemispheric correlations

Prior reports have suggested that the degree of volume correspondence of a given region across hemispheres within an individual is moderate to strong (Allen et al., 2002). As such, one metric that has been proposed to capture reliability within method (Schoemaker et al., 2016) quantifies the degree to which a given structure’s size in one hemisphere predicts its size in the other. A weak correlation between hemispheres for a given method could potentially reflect error in volume estimation. Furthermore, a drop in interhemispheric correlation for an automated method relative to manual may be indicative of lower performance in the automated approach (i.e., to the extent that manual delineation is close to the true level of symmetry). We performed across-participant Pearson’s correlations of left and right hemisphere volumes (hereafter termed inter-hemisphere correlations, IHC) for each method (including Manual for reference) and each region.

### Statistical analyses

For all quantitative metrics (DSC, ICC, IHC, and Bias), we used a nonparametric approach to compute 95% confidence intervals and p values. We resampled participants with replacement across 1,000 iterations, each time computing the statistic(s) of interest. Inferences were made on the basis of these bootstrapped distributions. All reported statistics were derived using this approach.

For DSC, ICC, and IHC measures, child and adolescent groups were each compared in a pairwise fashion to the adult group, which was treated as the baseline. Specifically, we generated a bootstrapped distribution of the difference between the two groups, which we then compared to zero to generate p values. In the case of the Bias analysis, F statistics for the main effects and interaction were computed across iterations, and the bootstrapped distribution was compared to one. As the main goal of our paper was to compare the degree to which semi-automated segmentations correspond with those generated manually across ages groups within a single method, we do not report direct comparisons across methods within each age group. However, the reader is invited to reference the 95% confidence intervals reported in each table to determine where significant differences exist across methods (i.e., where the mean of one method falls outside the 95% confidence intervals of a second method). This approach is recommended to inform the selection of a method that yields the best performance, either across the whole age range reported here or within a particular age group of interest.

As our goal is to show where differences across groups *might* exist at a liberal threshold, we do not correct for multiple comparisons in our main analyses. We note throughout the text and tables which pairwise comparisons would survive Bonferroni correction, where we correct for the number of tests across subfields and age groups within method (ANTs and ANTs_ROI_: 12 tests, ASHS_S_: 10 tests, ASHS_C_: 8 tests).

### Follow-up analysis omitting template and atlas subjects

As a subset of our participants went into creating the group ANTs template (N=10) and ASHS atlas (N=27), it might be the case that those participants drive the measures computed across the whole group. To determine whether this was indeed the case, we repeated all analyses described above, this time completely omitting those participants who went into template and atlas generation. Importantly, we omitted all participants who went into either the ANTs template or ASHS atlas (N=27; all ANTs template participants were also used in the ASHS atlases) from the analyses. We note that these analyses are likely underpowered due to the large drop in N compared with our main analysis (reduced N=35 instead of 62); thus, a large increase in variance is to be expected, especially for correlation-based measures (ICC, IHC). For completeness, these results omitting template participants are included on all barcharts as grey dots and error bars representing the mean and 95% confidence intervals; and on bias and edge overlap plots as dashed lines plotting the regression line and group means, respectively.

## RESULTS

### Comparing automated segmentation methods with manual demarcation

Our primary goal was to determine whether the degree to which semi-automated methods corresponded to regions delineated by an expert human rater varies across age groups in a pediatric sample. We thus assessed spatial overlap, edge agreement, volume correspondence, and potential bias of each automated method—ANTs, ANTs_ROI_, ASHS_S_, and ASHS_C_—relative to Manual. Due to the poor intra-rater reliability of manual segmentations in CA_2,3_ and subiculum, we do not discuss these two regions here.

#### Spatial overlap (DSC)

The main results of the spatial overlap analysis are presented in **Table 1** and **Figure 2**. Overall, DSC values neared or exceeded the agreement typically achieved by two human raters (ranging roughly from 0.7-0.85 in Olsen et al., 2013; though specific values will be resolution-dependent, Yushkevich et al., 2015b). For all regions, the ASHS approaches yielded better validity than the ANTs methods; comparing the two ANTs approaches, ANTs_ROI_ outperformed ANTs in all cases, suggesting that landmark-guided normalization may be superior when aiming to achieve maximal spatial overlap. Age-related differences were assessed by comparing child and adolescent groups to the adults using nonparametric t-tests. There were small but reliable differences at a liberal threshold of uncorrected p<0.05 between age groups in overall HPC (children and adolescents for ANTs and ASHS_S_; children using ASHS_C_) and DG/CA_2,3_ (adolescents using ANTs). There were no differences for CA_1_ and DG. Only the difference between children and adults for HPC using both ASHS methods remained significant after correcting for multiple comparisons; moreover, the differences in overlap values are numerically quite small (0.01-0.04) and are lessened by using ASHS_C_ (all < 0.02). Spatial overlap within the HPC head and body subfields separately revealed a slight advantage in agreement for subfields within the body over the head (**Inline Supplementary Tables S1-2** and **Inline Supplementary Figures S2-3**). Results were similar across left and right hemispheres (**Inline Supplementary Tables S3-4** and **Inline Supplementary Figures S4-5**).

**Table 1.** Spatial overlap of each method with Manual ROIs. Mean DSC and lower and upper bounds of 95% confidence intervals across all participants (left), as well as for child, adolescent, and adult groups separately. Asterisks indicate significance level of nonparametric t-tests comparing child and adolescent groups, respectively, with adults. * p < 0.05 and ** p < 0.01, uncorrected. ^+^ survives correction for multiple comparisons within method. Parentheses around SUB and CA_2,3_ indicate that these regions fell below our intra-rater reliability threshold (ICC(2,1)<0.80) and thus we do not consider them in the text. See also **Figure 2**.

| ROI | All<br>DSC [95%CI] | Child<br>DSC [95%CI] | Adolescent<br>DSC [95%CI] | Adult<br>DSC [95%CI] |
| --- | --- | --- | --- | --- |
| **ANTs** |  |  |  |  |
| HPC | 0.84 [0.83 0.85] | 0.83 [0.81 0.85]* | 0.84 [0.83 0.85]* | 0.86 [0.84 0.87] |
| CA<sub>1</sub> | 0.70 [0.69 0.70] | 0.69 [0.67 0.71] | 0.69 [0.68 0.71] | 0.70 [0.69 0.72] |
| (SUB) | 0.65 [0.64 0.66] | 0.65 [0.63 0.66] | 0.65 [0.63 0.66] | 0.67 [0.65 0.69] |
| DG/CA<sub>2,3</sub> | 0.78 [0.77 0.79] | 0.78 [0.76 0.79] | 0.78 [0.77 0.79]* | 0.79 [0.78 0.80] |
| (CA<sub>2,3</sub>) | 0.44 [0.42 0.45] | 0.42 [0.39 0.44]* | 0.44 [0.42 0.45] | 0.46 [0.43 0.48] |
| DG | 0.74 [0.73 0.74] | 0.73 [0.72 0.75] | 0.74 [0.73 0.75] | 0.74 [0.72 0.75] |
| **ANTs<sub>ROI</sub>** |  |  |  |  |
| HPC | 0.90 [0.90 0.90] | 0.90 [0.89 0.90] | 0.90 [0.89 0.90] | 0.90 [0.90 0.91] |
| CA<sub>1</sub> | 0.73 [0.72 0.74] | 0.73 [0.72 0.74] | 0.72 [0.71 0.73] | 0.73 [0.72 0.74] |
| (SUB) | 0.73 [0.72 0.73] | 0.73 [0.72 0.74] | 0.72 [0.71 0.73] | 0.73 [0.72 0.74] |
| DG/CA<sub>2,3</sub> | 0.81 [0.80 0.82] | 0.81 [0.80 0.82] | 0.80 [0.79 0.81] | 0.81 [0.81 0.82] |
| (CA<sub>2,3</sub>) | 0.48 [0.47 0.49] | 0.47 [0.45 0.49] | 0.48 [0.46 0.50] | 0.50 [0.48 0.51] |
| DG | 0.75 [0.74 0.75] | 0.75 [0.74 0.75] | 0.74 [0.74 0.75] | 0.75 [0.74 0.76] |
| **ASHs<sub>s</sub>** |  |  |  |  |
| HPC | 0.92 [0.92 0.93] | 0.92 [0.91 0.92]*** | 0.92 [0.91 0.93]* | 0.93 [0.93 0.93] |
| CA<sub>1</sub> | 0.80 [0.79 0.81] | 0.80 [0.79 0.81] | 0.81 [0.79 0.82] | 0.80 [0.79 0.81] |
| (SUB) | 0.78 [0.77 0.79] | 0.78 [0.76 0.79] | 0.77 [0.76 0.79] | 0.79 [0.78 0.80] |
| DG/CA<sub>2,3</sub> | -- | -- | -- | -- |
| (CA<sub>2,3</sub>) | 0.63 [0.62 0.65] | 0.61 [0.58 0.64]* | 0.64 [0.62 0.66] | 0.65 [0.63 0.68] |
| DG | 0.84 [0.83 0.84] | 0.84 [0.83 0.85] | 0.84 [0.83 0.85] | 0.84 [0.82 0.85] |
| **ASHs<sub>c</sub>** |  |  |  |  |
| HPC | 0.92 [0.92 0.93] | 0.92 [0.91 0.92]*** | 0.92 [0.92 0.93] | 0.93 [0.93 0.94] |
| CA<sub>1</sub> | 0.81 [0.80 0.82] | 0.80 [0.79 0.81] | 0.81 [0.80 0.83] | 0.81 [0.80 0.83] |
| (SUB) | 0.79 [0.78 0.80] | 0.78 [0.77 0.79]* | 0.78 [0.76 0.81] | 0.80 [0.79 0.82] |
| DG/CA<sub>2,3</sub> | 0.87 [0.87 0.88] | 0.87 [0.85 0.88] | 0.88 [0.86 0.89] | 0.88 [0.87 0.89] |
| (CA<sub>2,3</sub>) | -- | -- | -- | -- |
| DG | -- | -- | -- | -- |

**Figure 2.**
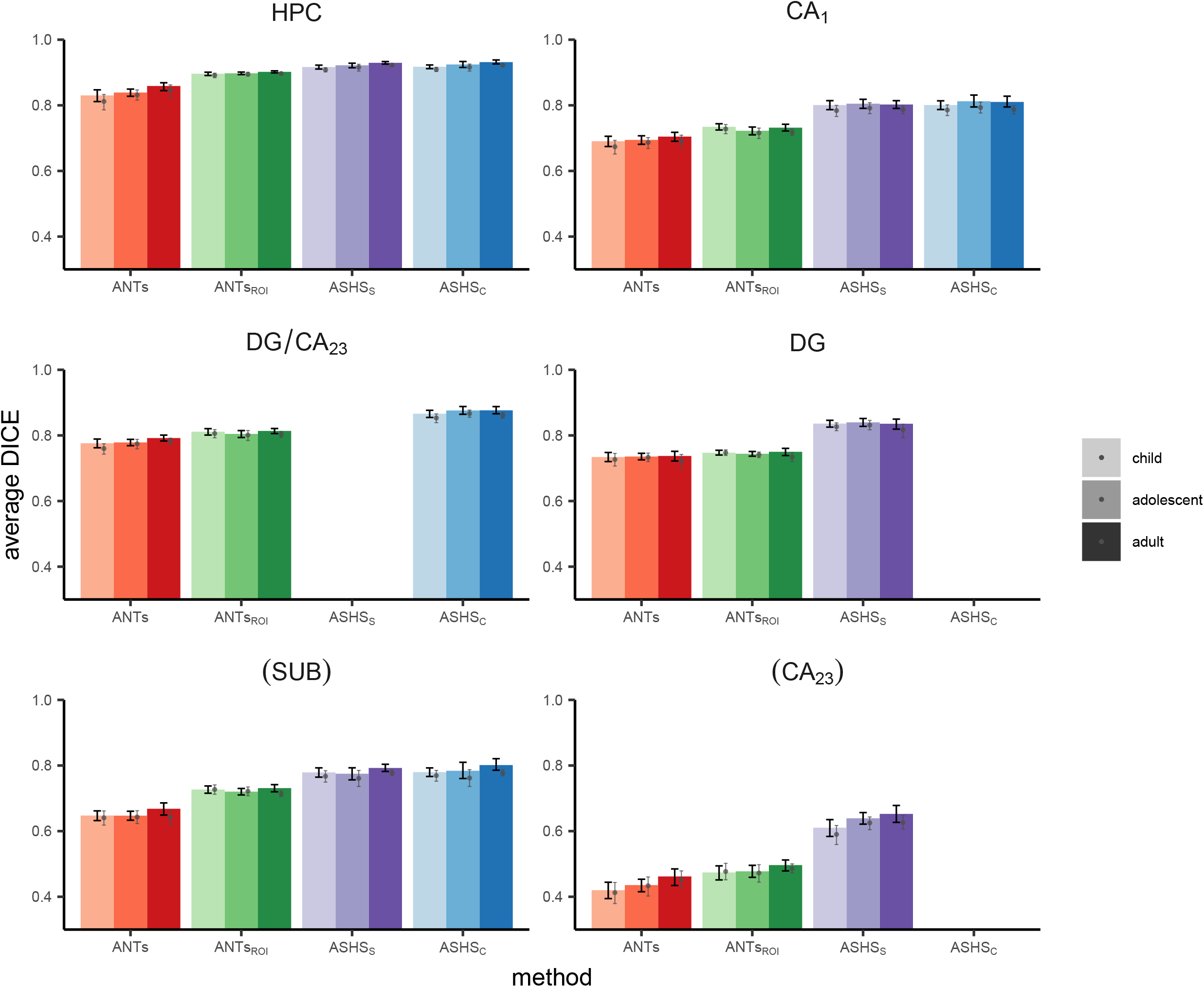
Spatial overlap of each method with Manual ROIs measured using DSC. Black error bars represent 95% confidence intervals on the main analysis. Grey dots and corresponding error bars represent means and 95% confidence intervals, respectively, for analysis omitting all N=27 participants who went into the generation of the ANTs template or ASHS atlases. Parentheses around SUB and CA_2,3_ indicate that these regions fell below our intra-rater reliability threshold (ICC(2,1)<0.80) and thus we do not consider them in the text. Data correspond with **Table 1**.

#### Edge agreement across methods

A voxelwise depiction of edge agreement is displayed in **Figure 3**. The edge between CA_1_ and DG, which largely relies on boundaries visible on MRI, were generally quite high across most of the anterior-posterior extent of the HPC, as well as across age groups. The outer edge of the overall HPC was also quite reliable across all methods, particularly on the lateral portions of the structure; there was relatively less agreement on how far DG/CA_2,3_ (particularly in the HPC head) and subiculum should extend in the medial and ventral directions, respectively. As would be expected, agreement was lower for those divisions lacking visible anatomical boundaries, such as between CA_1_ and subiculum and between CA_2,3_ and DG in the head. However, there was also substantial disagreement between methods in the stratum radiatum lacunosum moleculare (SRLM), which is somewhat surprising given that this is a highly visible boundary. It did seem to be the case that there was less overall agreement about where edges should be placed in the head relative to the body of the HPC. Despite these differences in agreement across the structure, there were no apparent differences in this pattern across age groups.

**Figure 3.**
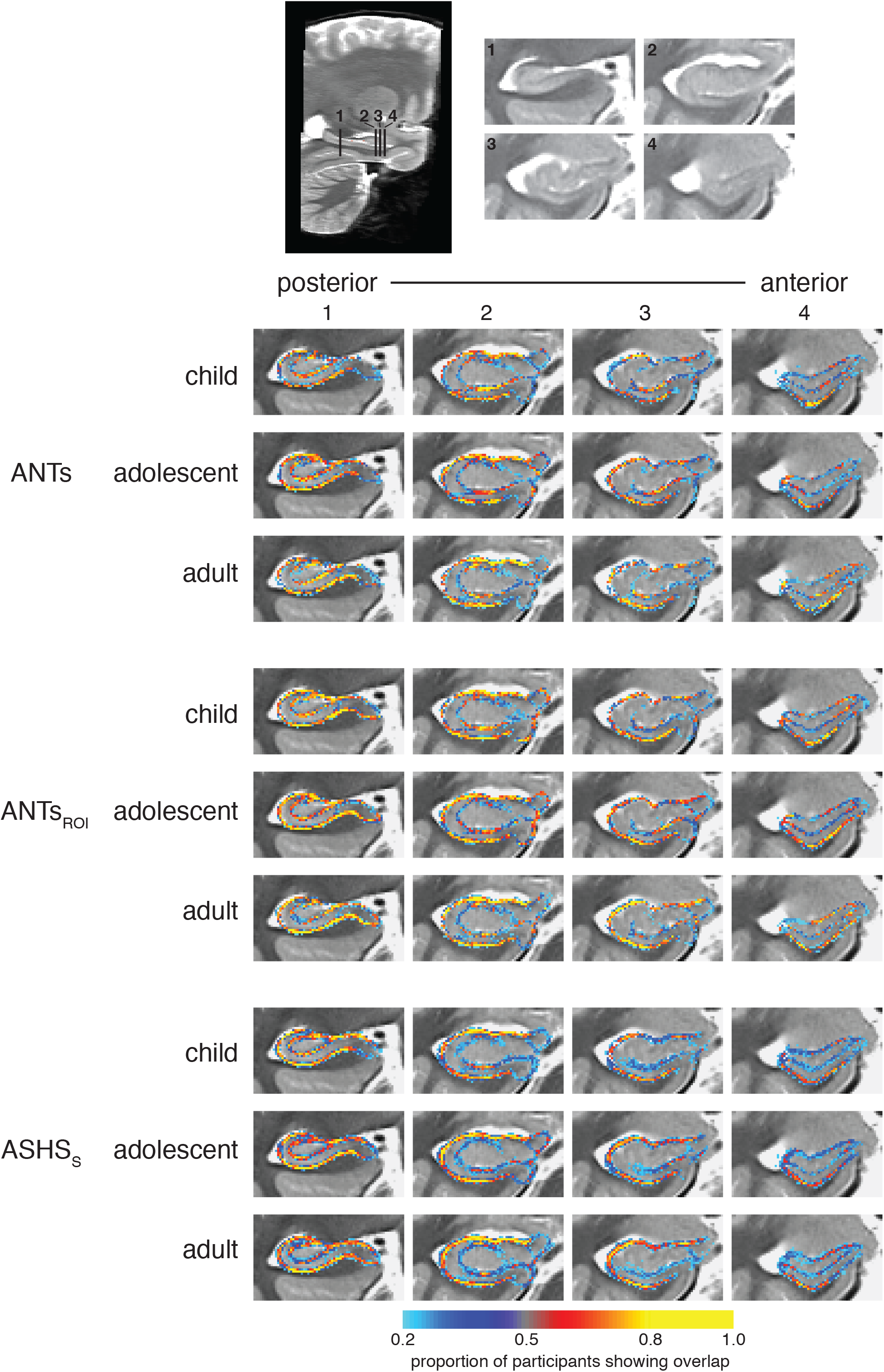
Voxelwise edge agreement displayed on a custom template separately for children, adolescents, and adults. Intensities represent the proportion of participants for which the method and Manual agreed that the voxel was a subfield boundary.

We next quantified overlap of these subfield edges as a function of position along the anterior-posterior axis of HPC to ask whether edge overlap differed between the HPC head versus body and tail. The slice-by-slice edge-based DSC results depicted in **Figure 4** show a consistent pattern across methods and age groups: there was generally higher agreement in the HPC body/tail relative to the head. By focusing on the edges rather than the entire volume of the subfields, this very stringent index showed a considerable advantage for ASHS over the both ANTs and ANTs_ROI_ methods. Of note, the degree of the advantage was not necessarily apparent from the standard DSC measures that consider the whole area of each subfield. In particular, the slice with the best spatial overlap for the ANTs methods is about the same as the slice with the worst overlap for ASHS. The adult group showed greater overlap than children and adolescents across most of the hippocampal axis. However, despite these differences, the child and adolescent agreement for ASHS was quite high overall and superior to edge overlap in adults using either ANTs method. Analyzing left and right hemispheres separately yielded similar results (**Inline Supplementary Figure S6**).

**Figure 4.**
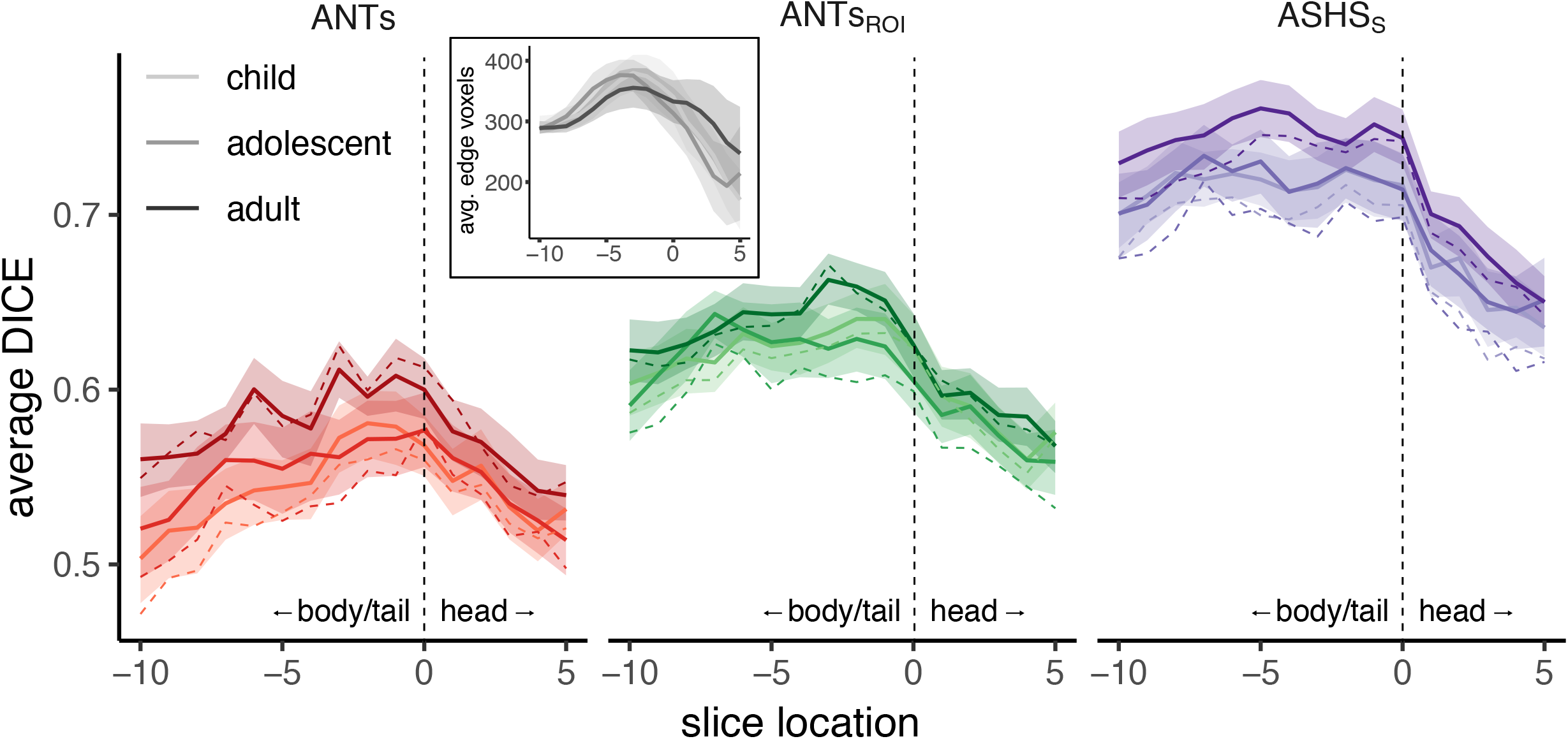
DSC edge overlap as a function of position along the anterior-posterior axis. Spatial overlap of edge maps for ANTs, ANTs_ROI_, and ASHS_S_ with Manual ROIs for each age group within left (top) and right (bottom) HPC. Lines represent group means; shaded regions represent 95% confidence intervals. Each participant’s hippocampus was centered on the slice dividing the head from the body, represented at zero with a dashed vertical line. Positive values along the x-axis (to the right of the dashed line) are in the HPC head; negative values (left) are in the remainder of HPC (a combined body/tail region). Overlap generally tracked with number of voxels going into the analysis (inset), which also varies as a function of anterior-posterior slice.

#### Volume correspondence: Intra-class correlation

In the next analysis, we investigated the degree to which each subfield volumes derived from the semi-automated methods corresponded with Manual, irrespective of their spatial agreement. These results are shown in **Table 2** and **Figure 5**. Comparing ICC measures, we found that most methods provided acceptable correspondence for overall HPC and CA_1_. A notable exception was ANTs, which demonstrated lower correspondence than the other methods in all ROIs for all three age groups (although showed no reliable differences as a function of age). Validity was lower and more variable in DG and DG/CA_2,3_ for all methods except ASHS_C_. It is worth noting that the ASHS methods yielded higher ICC values than either ANTs method almost universally across regions.

**Table 2.** Volume correspondence for each automated method with Manual ROIs. Mean ICC (absolute agreement of single measures) value and lower and upper bounds of 95% confidence intervals across all participants (left), as well as for child, adolescent, and adult groups separately. Asterisks indicate significance level of nonparametric t-tests comparing child and adolescent groups, respectively, with adults. * p < 0.05 and ** p < 0.01, uncorrected. No tests survived correction for multiple comparisons. Parentheses around SUB and CA_2,3_ indicate that these regions fell below our intra-rater reliability threshold (ICC(2,1)<0.80) and thus we do not consider them in the text. See also **Figure 4**.

| ROI | All<br>ICC [95%CI] | Child<br>ICC [95%CI] | Adolescent<br>ICC [95%CI] | Adult<br>ICC [95%CI] |
| --- | --- | --- | --- | --- |
| **ANTs** |  |  |  |  |
| HPC | 0.51 [0.35 0.63] | 0.37 [0.02 0.52] | 0.63 [0.34 0.78] | 0.56 [0.15 0.76] |
| CA<sub>1</sub> | 0.56 [0.39 0.67] | 0.47 [0.10 0.64] | 0.58 [0.26 0.76] | 0.63 [0.37 0.78] |
| (SUB) | 0.54 [0.36 0.70] | 0.29 [-0.08 0.55] | 0.70 [0.44 0.82] | 0.64 [0.34 0.91] |
| DG/CA<sub>2,3</sub> | 0.38 [0.26 0.46] | 0.39 [0.14 0.50] | 0.31 [0.12 0.45] | 0.40 [0.18 0.55] |
| (CA<sub>2,3</sub>) | 0.14 [0.07 0.20] | 0.15 [-0.01 0.25] | 0.12 [0.02 0.22] | 0.15 [0.03 0.25] |
| DG | 0.47 [0.31 0.57] | 0.49 [0.06 0.65] | 0.38 [0.08 0.57] | 0.49 [0.20 0.65] |
| **ANTs<sub>ROI</sub>** |  |  |  |  |
| HPC | 0.82 [0.76 0.86] | 0.84 [0.68 0.89] | 0.81 [0.67 0.89] | 0.83 [0.64 0.90] |
| CA<sub>1</sub> | 0.79 [0.68 0.85] | 0.79 [0.42 0.90] | 0.79 [0.60 0.88] | 0.79 [0.61 0.88] |
| (SUB) | 0.66 [0.50 0.78] | 0.66 [0.27 0.86] | 0.70 [0.48 0.83] | 0.61 [0.30 0.82] |
| DG/CA<sub>2,3</sub> | 0.47 [0.32 0.58] | 0.58 [0.40 0.71] | 0.38 [0.11 0.60] | 0.39 [0.02 0.60] |
| (CA<sub>2,3</sub>) | 0.16 [0.07 0.25] | 0.15 [-0.04 0.25] | 0.16 [0.01 0.32] | 0.18 [0.00 0.33] |
| DG | 0.47 [0.29 0.59] | 0.69 [0.35 0.78] | 0.33 [0.03 0.53] | 0.36 [-0.03 0.57] |
| **ASHS<sub>s</sub>** |  |  |  |  |
| HPC | 0.75 [0.64 0.82] | 0.71 [0.47 0.79]* | 0.67 [0.41 0.82]* | 0.89 [0.76 0.95] |
| CA<sub>1</sub> | 0.74 [0.63 0.82] | 0.70 [0.42 0.81] | 0.72 [0.48 0.86] | 0.81 [0.65 0.89] |
| (SUB) | 0.68 [0.54 0.78] | 0.65 [0.37 0.82] | 0.67 [0.38 0.81] | 0.73 [0.48 0.85] |
| DG/CA<sub>2,3</sub> | -- | -- | -- | -- |
| (CA<sub>2,3</sub>) | 0.26 [0.15 0.38] | 0.27 [0.10 0.45] | 0.22 [0.01 0.42] | 0.26 [0.04 0.43] |
| DG | 0.72 [0.57 0.81] | 0.77 [0.40 0.88] | 0.55 [0.24 0.80] | 0.81 [0.63 0.89] |
| **ASHS<sub>c</sub>** |  |  |  |  |
| HPC | 0.75 [0.65 0.82] | 0.71 [0.47 0.80]* | 0.68 [0.43 0.82]* | 0.89 [0.74 0.95] |
| CA<sub>1</sub> | 0.70 [0.59 0.78] | 0.65 [0.34 0.77] | 0.69 [0.46 0.84] | 0.78 [0.61 0.87] |
| (SUB) | 0.66 [0.52 0.77] | 0.62 [0.34 0.80] | 0.65 [0.34 0.81] | 0.74 [0.50 0.86] |
| DG/CA<sub>2,3</sub> | 0.72 [0.61 0.78] | 0.70 [0.40 0.79] | 0.63 [0.40 0.78] | 0.80 [0.66 0.87] |
| (CA<sub>2,3</sub>) | -- | -- | -- | -- |
| DG | -- | -- | -- | -- |

**Figure 5.**
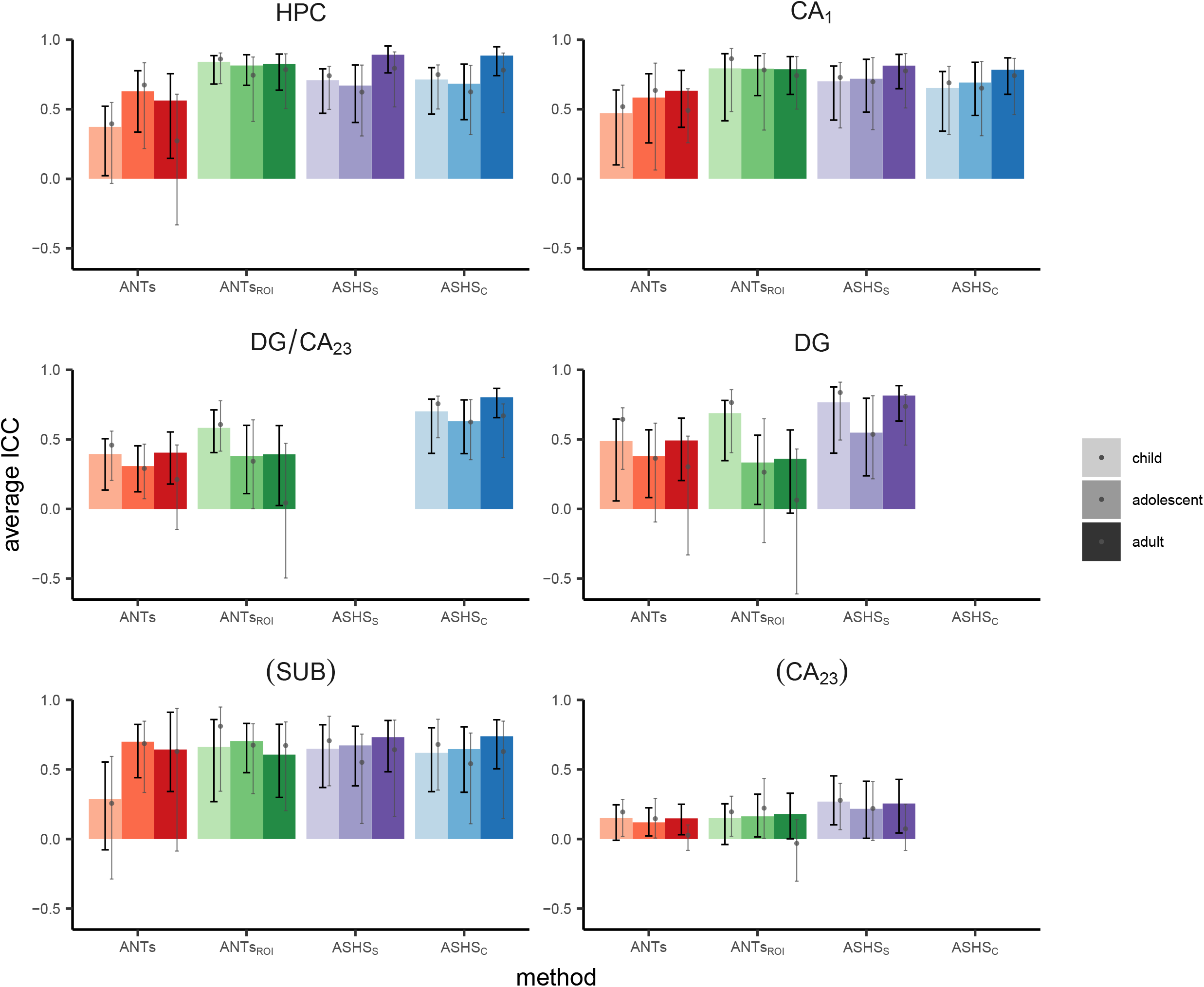
Volume correspondence of each method with Manual ROIs measured using ICC. Black error bars represent 95% confidence intervals on the main analysis. Grey dots and corresponding error bars represent means and 95% confidence intervals, respectively, for analysis omitting all N=27 participants who went into the generation of the ANTs template or ASHS atlases. Parentheses around SUB and CA_2,3_ indicate that these regions fell below our intra-rater reliability threshold (ICC(2,1)<0.80) and thus we do not consider them in the text. Data correspond with **Table 2**.

Echoing the DSC results, age-related differences were limited. One caveat to this result is that in some methods and regions ICC was quite low and not reliably different from zero; thus, a lack of group differences in such cases is difficult to interpret. We observed significantly better volume agreement in the adults relative to both the child and adolescent groups in overall HPC when using ASHS_S_ or ASHS_C_. However, these differences did not survive correction for multiple comparisons. There were no differences using any method in any of the subfields. In sum, ASHS_C_ provided the highest validity across methods within the tested ROIs (HPC, CA_1_, SUB, DG/CA_2,3_), with minimal evidence for age-related differences.

Dividing HPC into head and body showed a difference across ANTs and ASHS methods, with ASHS generally performing better in the body and ANTs generally performing better in the head (see **Inline Supplementary Tables S5-6** and **Figures S7-8**). However, given that ICC in the body for ANTs were in some cases not reliably different from zero, again these results should be interpreted with caution. See **Inline Supplementary Tables S7-8** and **Figures S9-10** for results split by hemisphere.

#### Bias

We generated Bland-Altman plots to look for potential bias in the volumes estimated by each automated method, shown in **Figures 6**. These plots show each automated method-Manual segmentation difference as a function of the average volume estimated across the two methods being compared. First, none of the automated methods showed evidence of overall bias as evidenced by confidence bands (dashed lines in Figure 6) encompassing zero in all cases. This is taken as evidence that none of the methods systematically over- or under-estimate the volumes relative to Manual.

**Figure 6.**
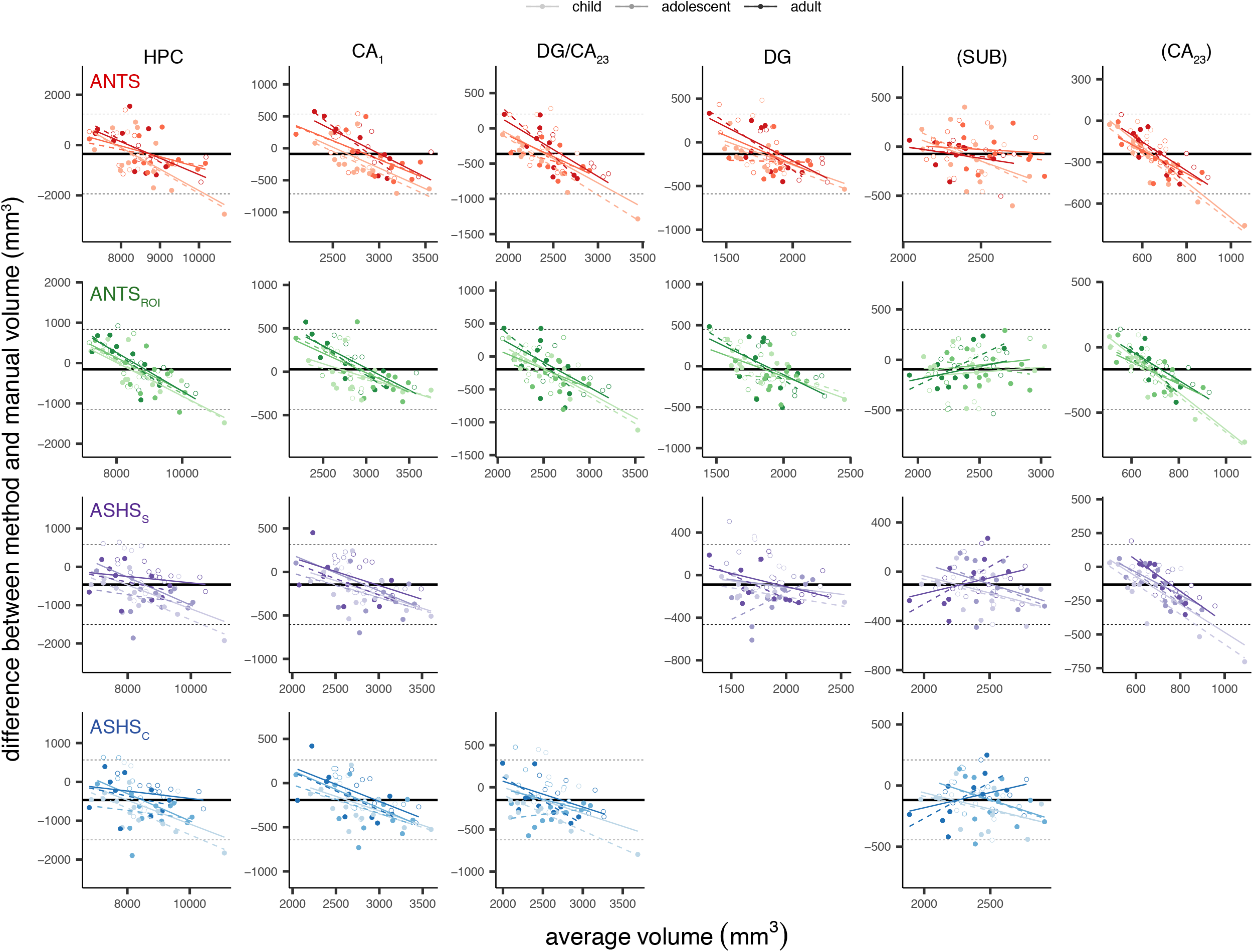
Bland-Altman plots comparing each automated method with Manual. Rows correspond with results from different automated methods, columns correspond with data from different ROIs. Within each plot, x-axis represents the mean regional volume across the two methods; y-axis represents the difference (method-Manual). Solid black line indicates the mean difference across all age groups; dashed lines are 2 standard deviations above and below the mean. Regression lines are displayed for each age group separately (dashed lines represent regression lines excluding atlas subjects). Parentheses around SUB and CA_2,3_ indicate that these regions fell below our intra-rater reliability threshold (ICC(2,1)<0.80) and thus we do not consider them in the text. ANCOVA statistics are provided in **Table 3**.

Second, we performed a series of ANCOVAs to test how the average volume, the age group, or the volume x age group interaction impacted the degree of discrepancy we observed between the two methods (**Table 3**). All regions we consider showed significant main effects of volume for all four methods; this indicates that the larger a region is, the more its volume tends to be underestimated by the automated methods. Significant main effects of age (p<0.05, uncorrected) were found only for CA_1_ using ANTs. These results suggest that the degree of bias differs reliably between the three groups only in this region using those specific methods. However, means split by age group were near zero in all cases, suggesting that despite these small differences all groups remain within the acceptable range. Of note, main effects of volume would survive correction for multiple comparisons for only a subset of regions. Splitting subfields into the HPC head and body revealed age-related bias only for ANTs and ANTs_ROI_ (see **Inline Supplementary Tables S9-10** and **Figures S11-12**). Age-related effects appeared to be slightly stronger in the right than left hemisphere (**Inline Supplementary Tables S11-12** and **Figures S13-14**). Taken together, these results suggest that while present in a few cases, the effects of age group on the degree of bias observed pale in comparison to the effect of region size. This represents an important bias that is pervasive in these methods (see Discussion for further consideration of this issue).

**Table 3.** Bias statistics for each automated method, with Manual ROIs serving as the baseline. F statistics and corresponding p values indicate the reliability of the main effects of volume, age, and volume x age interactions. * p < 0.05 and ** p < 0.01, uncorrected. ^+^ survives correction for multiple comparisons within method. Parentheses around SUB and CA_2,3_ indicate that these regions fell below our intra-rater reliability threshold (ICC(2,1)<0.80) and thus we do not consider them in the text. These data are depicted in **Figures 5-8**.

| ROI | Volume<br>F statistic (p value) | Age<br>F statistic (p value) | Volume x Age<br>F statistic (p value) |
| --- | --- | --- | --- |
| **ANTs** |  |  |  |
| HPC | 30.99 (<0.001)<sup>***</sup> | 1.52 (0.26) | 1.23 (0.34) |
| CA<sub>1</sub> | 39.79 (<0.001)<sup>***</sup> | 4.66 (0.02)* | 0.18 (0.66) |
| (SUB) | 4.00 (0.13) | 1.41 (0.27) | 1.09 (0.37) |
| DG/CA<sub>2,3</sub> | 50.19 (<0.001)<sup>***</sup> | 1.72 (0.21) | 0.22 (0.60) |
| (CA<sub>2,3</sub>) | 120.65 (<0.001)<sup>***</sup> | 2.20 (0.15) | 1.60 (0.24) |
| DG | 28.59 (<0.001)<sup>***</sup> | 0.97 (0.34) | 0.33 (0.57) |
| **ANTs<sub>ROI</sub>** |  |  |  |
| HPC | 62.20 (<0.001)<sup>***</sup> | 1.74 (0.19) | 0.15 (0.71) |
| CA<sub>1</sub> | 38.88 (<0.001)<sup>***</sup> | 2.64 (0.12) | 0.64 (0.53) |
| (SUB) | 1.77 (0.34) | 0.57 (0.43) | 0.16 (0.59) |
| DG/CA<sub>2,3</sub> | 41.16 (<0.001)<sup>***</sup> | 0.82 (0.39) | 0.25 (0.60) |
| (CA<sub>2,3</sub>) | 113.64 (<0.001)<sup>***</sup> | 2.65 (0.11) | 2.23 (0.17) |
| DG | 23.39 (0.000)<sup>***</sup> | 0.21 (0.54) | 0.23 (0.60) |
| **ASHS<sub>s</sub>** |  |  |  |
| HPC | 12.90 (0.01)* | 2.60 (0.11) | 1.63 (0.23) |
| CA<sub>1</sub> | 21.72 (<0.001)<sup>***</sup> | 1.38 (0.25) | 0.17 (0.66) |
| (SUB) | 1.40 (0.40) | 1.25 (0.25) | 3.95 (0.05)* |
| DG/CA<sub>2,3</sub> | -- | -- | -- |
| (CA<sub>2,3</sub>) | 74.60 (<0.001)<sup>***</sup> | 5.11 (0.04)* | 0.41 (0.52) |
| DG | 3.23 (0.19) | 0.09 (0.60) | 0.17 (0.57) |
| **ASHS<sub>c</sub>** |  |  |  |
| HPC | 13.43 (0.01)<sup>**</sup> | 2.76 (0.11) | 1.41 (0.28) |
| CA<sub>1</sub> | 28.79 (<0.001)<sup>***</sup> | 2.02 (0.16) | 0.14 (0.65) |
| (SUB) | 1.44 (0.40) | 1.63 (0.21) | 4.01 (0.05)* |
| DG/CA<sub>2,3</sub> | 11.07 (0.04)* | 0.51 (0.46) | 0.05 (0.57) |
| (CA<sub>2,3</sub>) | -- | -- | -- |
| DG | -- | -- | -- |

### Assessing reliability within segmentation method

#### Volume correspondence within an individual: inter-hemispheric correlations

In addition to characterizing how well each automated method corresponds with the Manual segmentation, we also wanted to quantify how consistent each method is with itself across participants’ two hippocampi. Previous work (Schoemaker et al., 2016) has investigated the correlation between volumes in the left and right hemispheres as an index of this within-method agreement, since individuals tend to have relatively similar structural volumes across hemispheres. We computed this measure for all methods including Manual, which serves as a baseline.

These data are displayed in **Table 4** and **Figure 7**. First, all regions considered here showed a significant relationship between left and right hemispheres across all methods. For the Manual ROIs, we observed no significant age group effects, indicating that the degree of correspondence across hemispheres changes little across development. However, when using the automated approaches, DG/CA_2,3_ for ANTs and ASHS_c_ showed significantly greater IHC for the child relative to the adult group. There were no group differences that survived correction for multiple comparisons. It seemed to be the case that the differences that emerged for the automated methods were driven by under- or overestimating the level of symmetry for a single group relative to the Manual regions, rather than a fundamental change in the pattern shown across all three groups. For instance, in DG/CA_2,3,_ three of the four automated methods yielded decreased IHC estimates relative to Manual in the adult group, while the estimates for the other age groups appear to remain unchanged. There were few differences in IHC between the HPC head and body with only slightly lower symmetry among adults in the body versus the head (see **Inline Supplementary Tables S13-14** and **Figures S15-16**).

**Table 4.** Within-method volume correspondence for all methods, indexed as Pearson correlations between left and right hemisphere volumes. Lower and upper bounds indicate 95% confidence intervals across all participants (left), as well as for child, adolescent, and adult groups separately. Note that the volume of CA_2,3_ is not reliably correlated across hemispheres in adults, making it difficult to interpret deviations from that group as a baseline. Asterisks indicate significance level of nonparametric t-test comparing child and adolescent groups, respectively, with adults. * p < 0.05 and ** p < 0.01, uncorrected. No tests survived correction for multiple comparisons. Parentheses around SUB and CA_2,3_ indicate that these regions fell below our intra-rater reliability threshold (ICC(2,1)<0.80) and thus we do not consider them in the text. See also **Figure 9**.

| ROI | All | Child | Adolescent | Adult |
| --- | --- | --- | --- | --- |
| **Manual** |  |  |  |  |
| HPC | 0.94 [0.90 0.96] | 0.95 [0.86 0.98] | 0.91 [0.84 0.97] | 0.95 [0.89 0.98] |
| CA<sub>1</sub> | 0.90 [0.85 0.94] | 0.85 [0.67 0.94] | 0.91 [0.85 0.96] | 0.94 [0.88 0.98] |
| (SUB) | 0.58 [0.41 0.72] | 0.67 [0.35 0.87] | 0.52 [0.22 0.80] | 0.55 [0.17 0.79] |
| DG/CA<sub>2,3</sub> | 0.79 [0.66 0.88] | 0.85 [0.64 0.95] | 0.74 [0.48 0.89] | 0.75 [0.48 0.90] |
| (CA<sub>2,3</sub>) | 0.70 [0.55 0.81] | 0.79 [0.52 0.92] | 0.55 [0.19 0.79] | 0.65 [0.46 0.81] |
| DG | 0.78 [0.66 0.87] | 0.84 [0.58 0.94] | 0.83 [0.53 0.93] | 0.72 [0.44 0.89] |
| **ANTs** |  |  |  |  |
| HPC | 0.80 [0.69 0.88] | 0.67 [0.38 0.89] | 0.85 [0.70 0.93] | 0.86 [0.67 0.95] |
| CA<sub>1</sub> | 0.73 [0.56 0.85] | 0.70 [0.48 0.87] | 0.76 [0.47 0.91] | 0.71 [0.25 0.92] |
| (SUB) | 0.45 [0.26 0.62] | 0.55 [0.27 0.74] | 0.35 [0.00 0.67] | 0.44 [0.12 0.72] |
| DG/CA<sub>2,3</sub> | 0.72 [0.60 0.82] | 0.87 [0.71 0.95]* | 0.72 [0.55 0.84] | 0.56 [0.11 0.80] |
| (CA<sub>2,3</sub>) | 0.45 [0.21 0.65] | 0.74 [0.44 0.92] | 0.33 [-0.16 0.68] | 0.28 [-0.26 0.64] |
| DG | 0.78 [0.68 0.86] | 0.85 [0.67 0.95] | 0.80 [0.59 0.92] | 0.69 [0.40 0.86] |
| **ANTs<sub>ROI</sub>** |  |  |  |  |
| HPC | 0.88 [0.81 0.92] | 0.92 [0.75 0.97] | 0.84 [0.72 0.93] | 0.87 [0.67 0.95] |
| CA<sub>1</sub> | 0.83 [0.73 0.90] | 0.80 [0.55 0.95] | 0.84 [0.71 0.92] | 0.87 [0.67 0.94] |
| (SUB) | 0.66 [0.51 0.79] | 0.70 [0.31 0.88] | 0.59 [0.26 0.83] | 0.73 [0.54 0.87] |
| DG/CA<sub>2,3</sub> | 0.65 [0.48 0.79] | 0.79 [0.55 0.91] | 0.67 [0.37 0.89] | 0.58 [0.20 0.84] |
| (CA<sub>2,3</sub>) | 0.51 [0.30 0.70] | 0.61 [0.32 0.84] | 0.62 [0.29 0.84] | 0.36 [-0.10 0.72] |
| DG | 0.69 [0.53 0.81] | 0.77 [0.45 0.91] | 0.72 [0.42 0.91] | 0.63 [0.27 0.85] |
| **ASHS<sub>s</sub>** |  |  |  |  |
| HPC | 0.90 [0.84 0.94] | 0.91 [0.81 0.97] | 0.88 [0.78 0.95] | 0.90 [0.76 0.96] |
| CA<sub>1</sub> | 0.79 [0.69 0.87] | 0.82 [0.63 0.93] | 0.79 [0.63 0.89] | 0.81 [0.60 0.93] |
| (SUB) | 0.70 [0.54 0.81] | 0.70 [0.38 0.87]* | 0.41 [0.04 0.67]** | 0.90 [0.80 0.96] |
| DG/CA<sub>2,3</sub> | -- | -- | -- | -- |
| (CA<sub>2,3</sub>) | 0.57 [0.37 0.72] | 0.76 [0.45 0.93]** | 0.54 [0.34 0.74]* | 0.06 [-0.49 0.47] |
| DG | 0.83 [0.75 0.90] | 0.89 [0.84 0.95] | 0.87 [0.74 0.94] | 0.78 [0.55 0.91] |
| **ASHS<sub>c</sub>** |  |  |  |  |
| HPC | 0.90 [0.85 0.94] | 0.93 [0.82 0.97] | 0.88 [0.76 0.95] | 0.90 [0.76 0.96] |
| CA<sub>1</sub> | 0.77 [0.66 0.85] | 0.81 [0.53 0.93] | 0.79 [0.62 0.90] | 0.76 [0.47 0.90] |
| (SUB) | 0.73 [0.58 0.83] | 0.76 [0.47 0.90] | 0.44 [0.09 0.70]** | 0.91 [0.83 0.96] |
| DG/CA<sub>2,3</sub> | 0.80 [0.72 0.87] | 0.90 [0.84 0.95]** | 0.84 [0.71 0.92] | 0.67 [0.41 0.84] |
| (CA<sub>2,3</sub>) | -- | -- | -- | -- |
| DG | -- | -- | -- | -- |

**Figure 7.**
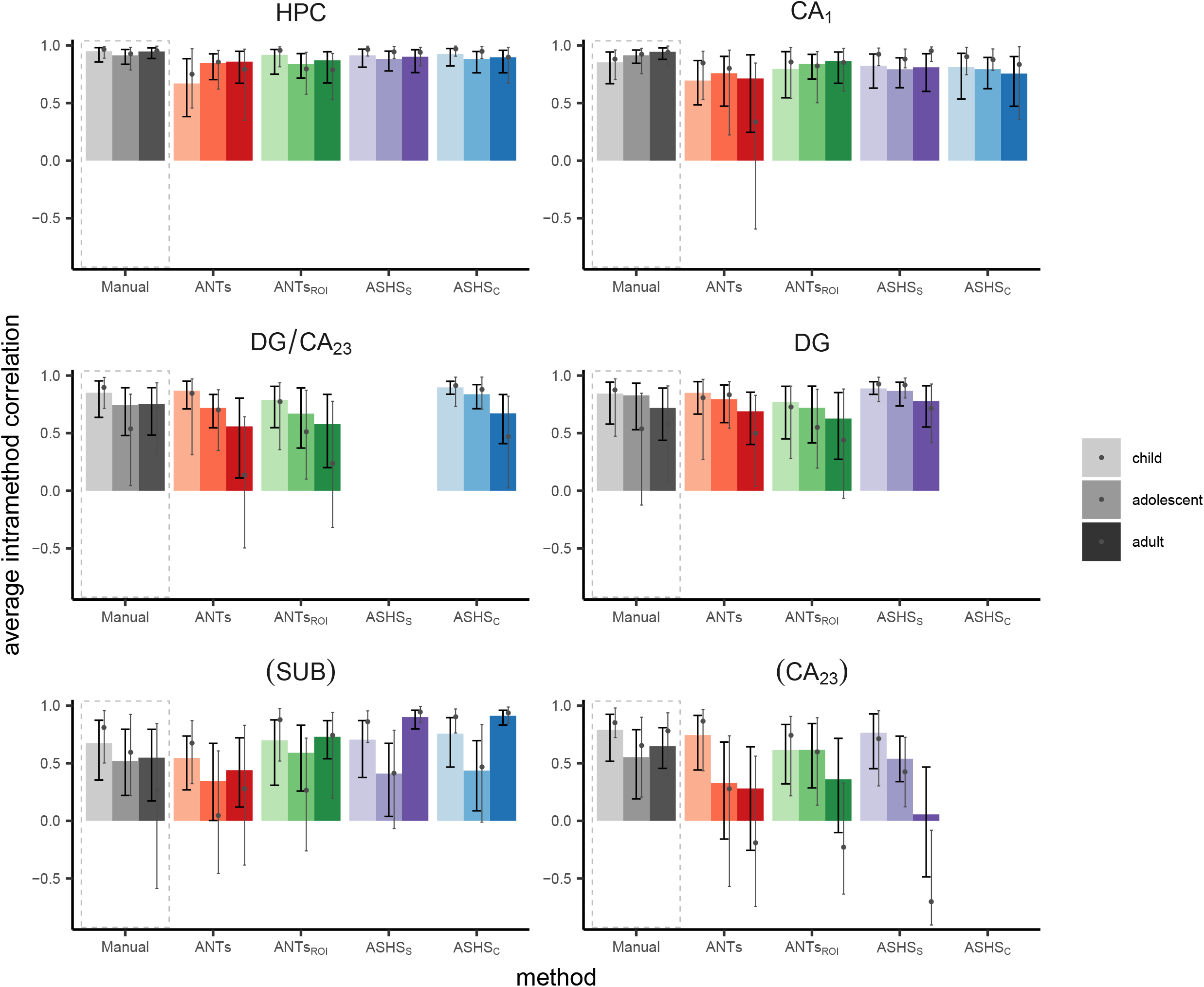
Within-method reliability assessed using IHC. Bar height represents the Pearson’s r value between the left and right hemisphere volumes for each group separately; black error bars represent 95% confidence intervals on the main analysis. Grey dots and corresponding error bars represent means and 95% confidence intervals, respectively, for analysis omitting all N=27 participants who went into the generation of the ANTs template or ASHS atlases. Parentheses around SUB and CA_2,3_ indicate that these regions fell below our intra-rater reliability threshold (ICC(2,1)<0.80) and thus we do not consider them in the text. Data correspond with **Table 4**.

### Impact of template/atlas participant omission

To assess the generalizability of our findings, we also calculated all metrics reported here omitting all N=27 participants who went into either the ANTs template or ASHS atlas generation. These results are displayed on figures throughout as grey dots (means) and error bars (95% confidence intervals) on bar charts and dashed lines on scatterplots and line graphs (representing regression line and group means, respectively). It should be noted that omitting such a large number of participants leaves a very small number of participants in each age group (child N=13, adolescent N=12, adult N=10). As such, correlation-based metrics (ICC, IHC) are not expected to be accurate reflections of the underlying relationship and may at times yield extreme values (e.g., CA_2,3_ for IHC in adults; **Figure 7**).

Generally speaking, the analyses omitting template participants showed similar means but wider confidence intervals compared with those metrics derived from the full sample. The similarity of the means across these analyses suggest that the template participants do not have a disproportionate impact on the pattern of results we see here. These initial results also speak to the generalizability of our ANTs templates and ASHS atlases and suggest these resources may be successfully applied to datasets with similar imaging protocols and samples.

## DISCUSSION

We compared HPC subfield regions of interest defined using two semi-automated methodological approaches to those demarcated by an expert human rater. Participants were children, adolescents, and adults, spanning an age range of approximately 6 through 30 years.

We chose methods that allowed us the flexibility to match the tracing protocol to the one we implemented manually, removing the concern that differences in the atlases used across methods may somewhat artificially lower performance (de Flores et al., 2015; Schoemaker et al., 2016; Wisse et al., 2014). We found generally good convergent validity across the methods—especially ASHS—and few differences for children and adolescents relative to the adult comparison group. Our findings extend previous work assessing the validity of automated methods designed to segment overall HPC in pediatric samples (Guo et al., 2015; Schoemaker et al., 2016). While these prior studies have yielded mixed results, our data complement recent work by Bender and colleagues (Bender et al., 2018) in demonstrating that there do exist methods that will segment the HPC into subfields (in addition to overall HPC) in pediatric samples in a way that approximates manual segmentation by an expert human rater. Building upon this prior work (Bender et al., 2018), our results additionally quantify segmentation performance as a function of age group, demonstrating that age does not appear to be a major factor in the validity of subfield segmentation methods.

One combination of region and method that showed poor performance was DG and DG/CA_2,3_ using ANTs_ROI_. In particular, ICC was low in these regions, with confidence intervals nearing or crossing zero in many cases in the adult group. Performance was particularly poor in the hippocampal body (**Inline Supplementary Table S6**). ASHS performance was far superior for these regions. As these particular regions are often the main target of developmental memory research given their late development (Lavenex and Banta Lavenex, 2013), this finding is particularly significant. These results suggest that, for researchers wishing to target DG or DG/CA_2,3_ combined subfield, ASHS is the far better choice.

One interesting pattern that stands out in our IHC analysis is the trend for a general decrease in symmetry over the course of development, a finding particularly evident in the subfields. A caveat to these findings is that IHC is relatively low in adults for some methods, which could reflect either error or true asymmetry. As this trend is numerically present even in the Manual regions, we suggest that this most likely reflects true asymmetry in neuroanatomical structure rather than bias in the automated methods. These results could reflect that hemispheric specialization of HPC emerges over developmental time. In adults, lateralized effects are often observed in terms of HPC task-based activation (Addis et al., 2011; Glosser et al., 1995; Golby et al., 2001; Kelley et al., 1998; Mack and Preston, 2016; Martin et al., 1997; Schlichting et al., 2014; Zeithamova and Preston, 2010), suggesting that the two hippocampi may serve unique and specialized roles. It may be the case that the general decrease in symmetry we observed is related to emerging specialization, as the two hemispheres progress along distinct developmental trajectories. Consistent with this notion, prior work has shown greater developmental change in the right than left HPC overall (Dennison et al., 2013; c.f. Daugherty et al., 2016). Similar ideas have been proposed for developmental emergence of hemispheric specialization in the visual stream (Behrmann and Plaut, 2015; Dundas et al., 2013); whether developmental asymmetry similarly reflects shifts in functional specialization across hemispheres in HPC subfields as well remains to be tested in future studies.

### Strengths of automated methods

Previous reports have used automated methods to draw conclusions about HPC structure in pediatric populations (Guo et al., 2015; Krogsrud et al., 2014; Lin et al., 2013; Tamnes et al., 2014a). For instance, one study (Lin et al., 2013) used automated methods to characterize maturational differences in HPC shape among children from 6-10 years of age. Their approach yielded findings that were not detectable when comparing volume alone, demonstrating the utility of applying such analyses to developmental questions. The authors applied three different automated methods and found that they gave similar results, suggesting the convergent validity of these methods even in their young participants. However, to the best of our knowledge, only three reports have quantified the validity of automated segmentation methods to yield HPC volumes in developmental samples and compared them with manual tracing. One such study (Schoemaker et al., 2016) found that both Freesurfer and FSL-FIRST overestimated overall HPC volumes, suggesting that these methods are not good approximations of manual tracing. Importantly, however, this problem does not seem to be unique to the children that were participants in that study, as other papers in adults have shown low validity using these methods (Doring et al., 2011; Pardoe et al., 2009).

Another report used MAGeT-Brain to segment the hippocampi of preterm neonates (Guo et al., 2015). That paper showed that segmentations derived from the automated methods were comparable to those performed manually, suggesting that automated methods can (in their case) be used with even the youngest of participants. A recent study (Bender et al., 2018) found good correspondence between automated segmentation using ASHS and manual tracing of the hippocampal body among an early lifespan sample (6-24 years). Performance was highest when the atlas was optimized—namely, that study suggests that a more variable atlas sample (as we have utilized in the current study) yields superior segmentations. However, it is unknown whether manual-ASHS correspondence in that study varied systematically with age. Taken together, these mixed findings in the prior literature underscore that the performance of the particular automated method being used must be assessed, as no two automated methods are the same in all cases.

Our findings extend beyond prior work to show that there exist automated methods for HPC subfield segmentation that can be used reliably in pediatric populations. Although previous work evaluating automated methods have included pediatric subjects (e.g., Bender et al., 2018), no study to date has specifically characterized the validity of such methods separately for child, adolescent, and young adult age ranges. We showed minimal age-related differences in the convergent validity of the semi-automated methods we tested. Of course, one caveat to this finding is our relatively small sample size; it is possible that significant relationships would be observed in a much larger group of participants. Nevertheless, we believe these results are welcome news for several reasons. While manual tracing is considered the best available approach for subfield segmentation, it has undeniable drawbacks: it is a laborious, time-consuming process that requires extensive training and anatomical expertise; it requires subjective decisions to be made on the part of the rater, leading to variability both within and across individuals—even those trained on the same tracing protocol—and is susceptible to bias and error; and varies substantially across research groups (Paul A. Yushkevich et al., 2015a). These issues are each expounded upon in turn below.

With respect to training needed to carry out this procedure, it is relatively minimal. At the data collection stage, scanner operators need training to monitor data as it is incoming and learn to judge what makes an “acceptable” coronal scan. After that point, both automated methods described in this paper require relatively little hands-on involvement on the part of the researcher. For our experienced rater at the resolution described in this paper, manual delineation of subfields takes roughly two hours; for less experienced raters, the process will take much longer. Most of the human involvement in the automated methods described here comes at the template or atlas generation stage: the researcher must create an acceptable template or atlas (or select among those publicly available), which not only takes time but also introduces a degree of human subjectivity that would be expected to propagate through the analysis pipeline to the final segmentations. Put another way, semi-automated methods are not purely objective. However, automated methods do significantly reduce the segmentation burden after this stage: once a template or atlas is generated, the ANTs approach we describe requires manual delineation on just one brain (the template), representing an investment of just a few hours; for ASHS, virtually no hands-on researcher time is needed. Depending on the resources available, these methods may require several hours of computation time to complete for a single participant; however, this is time during which the researcher can be doing other things. The knowledge of HPC subfield anatomy required is also quite limited. There are options available—such as using ASHS with an existing atlas—that require little anatomical expertise to carry out, and (in our experience) have a very low failure rate.

A clear strength of automated approaches is that they reduce the inconsistencies associated with human raters (Lerma-Usabiaga et al., 2016). Such inconsistencies may arise from simple differences in subjective tracing decisions across individuals (e.g., one rater tending to draw regions slightly more “generously” than another) or even within an individual, across occasions. Most relevant to the present report, knowledge of participant age might subconsciously influence raters to delineate regions in a certain way. While measures can be taken to avoid this issue in the case of development (e.g., by blinding the rater to subject identity and cropping the field of view to obscure identifying features like head size), this approach might be difficult or impossible in other special populations (e.g., those with brain damage or obvious atrophy). Thus, automated approaches that significantly reduce human subjectivity may be an ideal solution. Unlike other automated methods (Desikan et al., 2006), those we describe here—particularly ANTs_ROI_ and ASHS—do not require manual editing or painstaking examination of regions. While basic quality assessment should certainly be performed, failures with these methods tend to be extreme (though rare) and quite obvious, even to an untrained eye. This fact reduces not only the burden of anatomical knowledge required, but also alleviates issues associated with the potential bias introduced by human subjectivity, even compared with other automated methods.

The way manual tracing is implemented also varies substantially across research groups (Paul A. Yushkevich et al., 2015a), making it difficult to compare findings across studies. Both ANTs templates and ASHS atlases are small enough in terms of file size that they can be easily shared with other research groups and/or made freely available for download through online repositories. These methods can be readily applied to new datasets, thus enabling direct comparisons across studies—even those carried out by different research groups. Furthermore, as the field’s knowledge of hippocampal anatomy continues to be refined, new atlases can be generated and applied to existing datasets in a relatively straightforward and efficient manner.

In discussing these features, it is clear that automated segmentation methods align well with the aspirations expressed by so many in the field (e.g., Gorgolewski and Poldrack, 2016): science should be more reproducible and open. Moreover, in efforts to make findings more replicable in future studies, larger sample sizes are becoming the goal for some and the norm for many, particularly when individual differences are of interest. This makes manual delineation of hippocampal subfields an intractable approach for many researcher groups due to its labor-intensive nature. In developmental work specifically, this problem is compounded when individual differences vary by age, and/or when data is collected from the same large number of participants over multiple years. The automated methods described here would make hypotheses falling into these categories addressable in a way that is not only logistically feasible but also less susceptible to potential bias and subjectivity introduced by a human rater.

### Limitations and future considerations

While we purposefully chose to have a single rater perform all manual segmentations to reduce concerns about inter-rater reliability when generating our template and atlases, we recognize that this approach is not without limitations. Specifically, we were unable to determine the degree to which the automated segmentations generated here would correspond with *another* manual rater using the same protocol. Future work should strive to establish validity of the manual tracings through such comparison across expert raters. Thus, while our results provide quantification of the degree to which ANTs- and ASHS-based approaches can be used to generate new segmentations that approximate a given rater—and the relative performance of each method—it is not necessarily the case that they would correspond at the same level when tested across raters. Relatedly, we are not able to assess the degree to which the added variability of multiple human raters impacts (either positively or negatively) the semi-automated segmentation methods. All of these questions would be interesting avenues for future investigation.

Furthermore, one weakness of the ANTs methods in particular is that in this case, a single segmentation (by a single rater) on the group template serves as the basis for all individual participant segmentations. One could imagine an extension of our approach in which group template segmentations from multiple raters were somehow combined to create the template ROIs. Alternatively, it might be beneficial to segment the group template using ASHS rather than a human rater; in this case, the template image could simply be treated as another participant and the ROIs generated through the usual ASHS voting procedure. Such an approach could be useful if the researcher required their data to be in some common (i.e., template) space, and should be formally tested in future work.

Moreover, how the current results generalize to other brain regions, tracing protocols, and MRI acquisition protocols is also an important open question. Note that here, we consider only the HPC proper, and did not investigate subregions of MTL cortex. A formal test of these methods for MTL cortical segmentation in pediatric samples remains an open question for future studies. With respect to tracing protocols, it is possible that a similar comparison of automated methods using a different tracing protocol may lead to a different outcome. Thus, while our results serve as an important test to the flexibility of both software packages in generating reliable templates/atlases for hippocampal segmentation using a tracing protocol that was independent of this method’s development, future work will be needed to provide an empirical test of these questions of generalizability to different regions, raters, and protocols. It also remains to be tested whether manual tracing (e.g., by comparison of multiple raters on the same dataset) or semi-automated segmentation is more robust to differences in image quality, such as across images acquired using different MRI scanners and/or acquisition parameters. We did not systematically test different imaging parameters, and thus it is quite possible that a different MRI protocol—for example, perhaps by collecting a single, higher quality image rather than averaging two images— would improve performance of either manual tracers and/or semi-automated methods in a developmental context.

Another limitation of the present study is the poor intra-rater reliability of manual segmentations in CA_2,3_ and subiculum, which precluded assessment of the automated methods in these regions. While we report these results throughout the paper for completeness, the reader is cautioned against drawing any conclusions from the degree of correspondence between semi-automated and manual methods within these regions. In particular, if agreement is high, that could be an indication that the error inherent to the manual tracings is being reproduced by the semi-automated method(s). If agreement is low, this could either be due to the error in the original segmentations or weaknesses in the automated method. As such, future work with more reliably manual segmentations will be necessary to assess whether there are developmental differences in the performance of these semi-automated methods within CA_2,3_ and subiculum.

We were able to assess the generalizability of our ANTs templates and ASHS atlases to only those participants that did not go into generating the initial template/atlas. While the findings were encouraging in that the overall pattern of results held, the very small number of participants remaining ultimately led to noisier estimates across all metrics. As such, although these initial findings are encouraging for the general use of our generated templates and atlases, further validation of these resources with a much larger sample is needed.

The most stringent criterion for assessing segmentation methods is to require no differences of any kind across age groups. This criterion was not applicable in the present report; we do find some age-related differences. However, we suggest that the direction of these differences is important to bear in mind when thinking about how our results should inform future study. In particular, one must consider whether the direction of their developmental hypothesis would be confounded by the direction of differences observed here. For example, most empirical work and theoretical accounts in developmental cognitive neuroscience aim to understand the ways in which children are not yet like adults; relative to adult groups, children typically show lower performance, less differentiated neural signatures, lower data quality, and so on. We reason that in the majority of cases, the main concern would be if the current results showed automated segmentations were *worse* in children—such a finding would indicate that there is more error or variability in segmentation for the child group.

Our results show that evidence for this sort of confound is minimal. While there are a few group differences (children and/or adolescents showing worse segmentation than adults), they are either numerically small and in the context of extremely good performance (e.g., DSC for ASHS) or are statistically weak and do not survive correction for multiple comparisons (ICC, Bias). In fact, in some cases, the results appear to go in the opposite direction; for example, there was a general pattern of greater IHC among the younger groups relative to adults. Thus, given that the vast majority of targeted hypotheses in developmental cognitive neuroscience that expect worse performance or more errors in children, we believe that the limited differences observed in the current study are encouraging for semi-automated methods.

One strong pattern did in fact emerge across all methods tested, and nearly all regions: the degree of bias (i.e., over- or under-estimation of region size) was significantly related to the size of the region itself. In the majority of cases, this was borne out as greater error—that is, a larger difference between volumes derived from the target method versus those derived manually—for larger regions. Others have reported similar observations, albeit to varying degrees, in other datasets (Bender et al., 2018; de Flores et al., 2015; Guo et al., 2015; Schoemaker et al., 2016; Yushkevich et al., 2010; P.A. Yushkevich et al., 2015). One possibility is that this bias is an artifact of the method, whereby a raw difference of 500 mm^3^ means something different for a region that is 3500 versus 2500 mm^3^. Nevertheless, it is important to keep in mind that while the degree of over- or under-estimation of volume did not vary systematically across age groups, it did vary as a function of region size. This result was true even for ASHS, our best performing method. One unwanted consequence of this finding is that the range of volume values across participants is artificially compressed—that is, the maximum volume is effectively reduced due to the systematic under-estimation in this range. Such bias could potentially make it difficult to detect true relationships between volume and another variable of interest (e.g., behavioral performance).

Like other studies using these semi-automated methods (Bender et al., 2018; Paul A. Yushkevich et al., 2015b), we found that performance varies markedly across subfields. We thus suggest that comparisons across age groups within subfield are the most meaningful, and direct comparisons across subfields should be performed and interpreted with caution. This recommendation is especially true when comparing subfields that demonstrate low reliability (e.g., CA_2,3_).

Finally, it is worth explicitly noting one key limitation that is common to all studies of hippocampal subfield segmentation in pediatric populations: our results rest on the assumption that hippocampal histology in children will be similar to that of adults. This assumption is made due to the absence of histological data specific to children. Future studies that directly compare histology with MRI scans in children will be necessary to clarify that this is indeed the case.

### Suggestions for the field

It is likely the case that no semi-automated segmentation method will ever be a perfect substitute for a human’s expert anatomical knowledge. However, we believe that the automated methods discussed here may be even better suited than manual tracing to answer the kinds of questions posed by developmental researchers, given the considerations about feasibility, subjectivity, and bias described above. We thus present the following recommendations for developmental researchers interested in using automated approaches to study HPC subfield anatomy or function. These ideas are described with the caveat that we did not examine all possible automated segmentation methods or all tracing protocols; recommendations are based on the comparison of just those we did test.

When at all possible, analyses should be done in each participant’s native space using ROIs produced by ASHS. Performance of ASHS was comparable to or better than ANTs methods in all metrics and regions. One strength that ASHS has over ANTs—and likely a reason why it performs so well—is that more information is used to perform the individual participant segmentations. That is, rather than a single manual segmentation performed by an individual rater being the basis of each participant’s ROIs (as is the case for the ANTs methods as implemented here), all atlas participants “vote” for each segmentation. Despite its notable strengths, there are situations in which we could imagine ASHS would be the less desirable option to the researcher. One drawback of ASHS compared with ANTs is the relatively larger number of participants required for generating the ASHS atlas. For the purposes of the methodological comparisons we perform here, the most stringent approach is to not include these participants in the final analysis (i.e., have these participants be a separate “training” dataset, ideally with the same imaging parameters as the target sample). However, future empirical work could include the atlas participants by either 1) using only their manually traced ROIs that were required for atlas generation for all subsequent analyses, 2) generating many ASHS atlases that exclude individual atlas subjects in a cross validation approach (Paul A. Yushkevich et al., 2015b), or 3) excluding atlas subjects votes during label fusion in segmenting their own ROIs, as we have done in the current work. Subsequent analyses performed with the ROIs could include not only analysis of volume but also function; for example, these subfields could serve as anatomical ROIs (after alignment and resampling to functional space) to investigate both univariate and multivariate functional activation.

When voxelwise, group level analyses are an absolute necessity, our findings suggest that generating a custom coronal template—importantly, including participants representative of the whole age range—may be an acceptable approach, although we suggest caution. Most notably, our data suggest that inferences regarding the DG/CA_2,3_ region would be particularly problematic in this situation, given the subpar performance of both ANTs-based methods on nearly all of our metrics. While it has been suggested that the use of the MNI template is appropriate for use with children as young as 7 (Burgund et al., 2002; Kang et al., 2003), this template is not appropriate for localizing activations to specific hippocampal subfields. Our study is the first that we know of to generate a single custom group ANTs template (and the second to generate an ASHS atlas; Bender et al., 2018) spanning a developmental sample of the kind that would be used in high-resolution imaging studies optimized for the medial temporal lobes. This single group template is an important feature of the present method, as it enables direct voxelwise comparisons across individuals of different ages. We found limited age-related differences in our metrics for both ANTs methods, suggesting that the ability to normalize to the group template does not substantially differ for children and adolescents relative to adults.

We have a slight preference for using ANTs_ROI_ over ANTs in this kind of analysis strategy, given that spatial overlap—the metric that, we reason, is most directly related to warping each subject’s functional data or statistical maps to the group template—as superior for the ROI-guided implementation of ANTs across all of the regions we examined. Notably, researchers could opt to use the overall HPC derived from ASHS rather than manually trace the region on each participant to use the ROI-guided normalization, substantially reducing the time burden associated with the ANTs_ROI_ method. Moreover, as mentioned previously, researchers might opt to perform segmentation on the ANTs group template in an automated fashion using ASHS rather than through manual delineation, which has the potential to further improve performance.

Finally, given that rater subjectivity is such a concern with manual approaches to HPC segmentation, we would recommend that an automated method like ANTs or ASHS is utilized as a subsidiary analysis even in studies that ultimately rely on manual tracing of all participants. Despite being the best available segmentation method, manual tracing is prone to biases and error, and there is no guarantee that all problems with tracings will be identified during quality control. Leveraging an automated method can aid in identifying such problems: manually defined ROIs that significantly diverge from automated segmentations raise the flag for a more in-depth review of the tracing. In this situation, an automated method serves as an extended approach to quality control for correcting biases and errors due to manual tracing.

### Conclusions

Recent years have seen an increase in the number of studies on HPC subfield function. It is additionally becoming clear that many interesting questions relating to hippocampal development remain to be answered—not only in terms of characterizing the typical developmental trajectory, but also to better understand how the structural development goes awry in less typical scenarios like significant early life stress (Teicher et al., 2003), childhood obesity (Chaddock et al., 2010), and neurodevelopmental disorders (Schumann et al., 2004). Our findings suggest that automated subfield segmentation techniques can be applied to healthy individuals ranging in age from 6-30 years. While the convergent validity of these methods to atypically developing samples remains to be explored, the present results show the promise in this avenue. The ability to readily apply such methods to a diverse sample may result in increased sensitivity for diagnosis, making these findings of relevance to basic researchers and clinicians alike.

## Acknowledgments

Many thanks to Jessica Church-Lang, Tammy Tran, and Amelia Wattenberger for assistance with participant recruitment, data collection, and helpful discussions. We also thank the Texas Advanced Computing Center (http://www.tacc.utexas.edu) at The University of Texas at Austin for providing critical computing resources.

